# Development of inducible genetic systems to finely tune efficacy of InhA-targeting anti-tuberculous drugs

**DOI:** 10.64898/2026.09.25.754513

**Authors:** Eliana Sepulveda Ramos, Maria Sofia Deschutter, Vanessa Point, Thibault Brunet, Janïs Laudouze, Wafaa Achache, Céline Crauste, Stéphane Canaan, Gabriela Gago, Mickaël Maziero, Pierre Santucci

**Affiliations:** Aix Marseille Univ, CNRS, LISM, IM2B, Marseille, France; Institute of Molecular and Cellular Biology of Rosario (IBR-CONICET-UNR), Rosario, Argentina; IHU Méditerranée Infection, Aix-Marseille Univ., France; IBMM, Univ Montpellier, CNRS, ENSCM, 34060 Montpellier, France

**Keywords:** Gene silencing, Therapeutic targets, Antimicrobial resistance, Prodrug activation, *inhA*

## Abstract

Tuberculosis (TB), caused by *Mycobacterium tuberculosis* (*Mtb*), remains a leading global health threat, especially due to multidrug-resistant strains. The first-line drug isoniazid (INH) and second-line drugs ethionamide (ETH) and prothionamide (PTH), are prodrugs that target the essential enoyl-ACP reductase InhA, a key enzyme involved in mycolic acid biosynthesis. Resistance to these drugs primarily arises from genetic mutations that inactivate the enzymatic function of their respective bioactivating enzymes. While the recent development of chemical boosters has shown promising results in restoring drug susceptibility in these mutants, genetic tools to dynamically modulate these pathways and fine-tune resistance could be extremely valuable to develop the next-generation of anti-InhA drugs. In this context, we leverage CRISPR interference (CRISPRi) to develop inducible and reversible genetic systems to fine-tune antibiotic susceptibility in *Mycobacterium smegmatis* (*Msmeg*) and *Mtb*. By using our CRISPRi-based approach we first generated an inducible *inhA* hypomorph, an essential target required for mycolic acid production and bacterial growth, therefore confirming that our method is fully functional. Then, we developed genetic tools to selectively target *katG*, *ethA*, and the transcriptional repressor *ethR*, in order to generate conditional hypomorphs with altered antibiotic susceptibility profiles. Antibiotic susceptibility assays confirmed that the conditional hypomorphs selectively displayed high-level resistance or hypervulnerability to INH or ETH upon CRISPRi induction. Altogether, this work establishes CRISPRi as a powerful genetic platform for selectively fine-tuning mycobacterial drug susceptibility, with potential applications in developing next-generation anti-TB therapies.

## Introduction

Tuberculosis (TB), caused by *Mycobacterium tuberculosis* (*Mtb*), remains a major global health threat. In its 2025 Global TB Report, the World Health Organization (WHO) reported 1.23 million deaths and estimated 10.7 million new cases in 2024, making TB the leading cause of death from a single infectious agent [1].

The current WHO-recommended treatment regimen for drug-susceptible TB includes a 6-month course of 4 distinct drugs including isoniazid (INH), rifampicin (RIF), ethambutol (EMB), and pyrazinamide (PZA). This standard “RHZE” regimen displays a treatment success rate approximating 90% globally [1]. Despite its effectiveness, the RHZE regimen faces several limitations including long duration and potential side effects that can lead to poor patient adherence, especially in resource-limited settings. Early discontinuation of treatment not only fails to cure TB but also promotes the emergence of drug-resistant strains. In that context, the emergence of rifampicin-resistant TB (RR-TB) and multidrug-resistant TB (MDR-TB; defined as resistant to both RIF and INH), accounted for nearly 400 000 cases in 2024 and has been identified as the major obstacle to control TB worldwide, and subsequently END-TB by 2035 [2].

INH, and thioamides antibiotics such as ethionamide (ETH) and prothionamide (PTH), are recommended by the WHO as first and second-line agents for treating susceptible and drug-resistant pulmonary TB, respectively [3]. These three compounds are prodrugs that inhibit the NADH-dependent enoyl-ACP reductase InhA, a key enzyme that belongs to the FAS-II system and required for the synthesis of mycolic acids [4–7]. By blocking InhA, they disrupt cell wall integrity, leading to rapid bacterial death [8, 9]. Hence, in the HRZE regimen, INH constitutes a cornerstone anti-TB drug, mainly for its unique bactericidal ability against actively replicating *Mtb*, which reduces the bacterial load quickly and helps preventing disease progression and transmission [10].

Resistance to INH and alternative InhA-targeting drugs such as ETH and PTH, is primarily driven by loss of function mutations in the genes encoding their activating enzymes, *katG* (Rv1908c), *ethA* (Rv3854c) or *mymA* (Rv3083) [3, 11–13]. Additional resistance mechanisms have been also identified within the InhA-encoding sequence resulting in the well-established I194T amino-acid substitution that reduces binding with the NADH cofactor which is known for being essential for INH sensitivity [14]. Mutations within *inhA* (Rv1484) or *fabG1/mabA* (Rv1483) promoter regions were also reported. Both mechanisms are often associated with low-level INH resistance and ETH/PTH cross-resistance [6].

Over the last decades, several studies have identified very promising lead compounds that might prevent or revert these INH or ETH/PTH-associated resistance mechanisms. Willand and colleagues demonstrated that chemical targeting of the TetR-like *ethA* repressor EthR [13, 15–17] induces a conformational change that impairs its binding to the DNA operator, thereby increasing *ethA* transcription and resulting in greater ETH bioactivation and antibacterial activity [18]. Further studies have exploited this unique concept to boost bioactivation pathways, and therefore reduce the amount of drug required to sterilize bacterial load *in vitro* and *in vivo* but also circumvent resistance [19, 20]. Accordingly, the lead compound Alpibectir (Alp, BVL-GSK098) is now a strong clinical candidate as ETH booster [21–23]. This compound underwent successful short-term clinical trial (NCT06748937) and will require now longer clinical trials to assess its use as part of alternative regimens [23].

Similarly, the Stallings’ group has identified a small molecule called C10, that prevents the selection for INH-resistant mutants and restores INH sensitivity in INH-resistant *Mtb* strains harboring loss of function mutations in *katG*. Additional work revealed that C10 sensitizes *katG*-hypomorphs without enhancing the bioactivation of INH, by mainly acting on carbon flux and energy metabolism downstream of INH-NAD synthesis [24–26].

Altogether, these results confirm that InhA is one of the most clinically validated drug targets that still stands as a very attractive therapeutic target for both existing and novel anti-TB agents. Indeed, the development of direct InhA inhibitors, which do not require activation by KatG, EthA or MymA [27–29] or chemical boosters that limit resistance and/or sensitize *Mtb* to sub-lethal doses, are being actively pursued as a novel strategy to overcome resistance and improve treatment outcomes.

In this work, we capitalized from the well-established CRISPRi system [30–32] to develop a robust and inducible approach for tuning drug resistance and susceptibility in *Msmeg* and *Mtb*. By performing targeted-CRISPRi silencing of *inhA*, *katG*, *ethA* and *ethR*, we confirmed that these genes are critical for the action or bioactivation of the anti-tuberculosis prodrugs INH and ETH/PTH. By using chemically-induced genetic repression or synthetic constitutive overexpression, coupled with antibiotic susceptibility profiling assays, we demonstrate that genetic targeting of *katG*, *ethA* or *ethR*, enable to tune drug susceptibility, ranging from hyper-vulnerable to drug resistant phenotypes. Validation of such approaches and findings in *Mtb* confirmed that our experimental tools could serve as a robust platform for future applications in the drug discovery space, including the screening and development of direct InhA inhibitors or chemical boosters.

## Results

### CRISPRi-mediated targeting the mycobacterial FAS-II system by silencing the expression of the essential gene *inhA*

We recently demonstrated that our fluorescence-based plasmid constructs pJL32 and pJL34, derived from the pIRL series [31, 32], enabled an efficient targeted repression of genes of interest and fluorescence imaging [33]. These include essential genes such as *rpoB* or *mmpL3*, but also genes involved in cell-wall biosynthesis/remodeling such as the glycopeptidolipid translocase *mmpL4b* in both *Msmeg* and *Mycobacterium abscessus* model species [33, 34]. By capitalizing from these tools, we designed and generated two integrative pJL32 CRISPRi-based constructs, named pEAR14 and pEAR15, that harbor two distinct sequences encoding single-guides RNA (sgRNA) to target the expression of *inhA* (*MSMEG*_3151) within *Msmeg* FAS-II system (Fig. 1A; Fig. S1).

**Figure 1:**
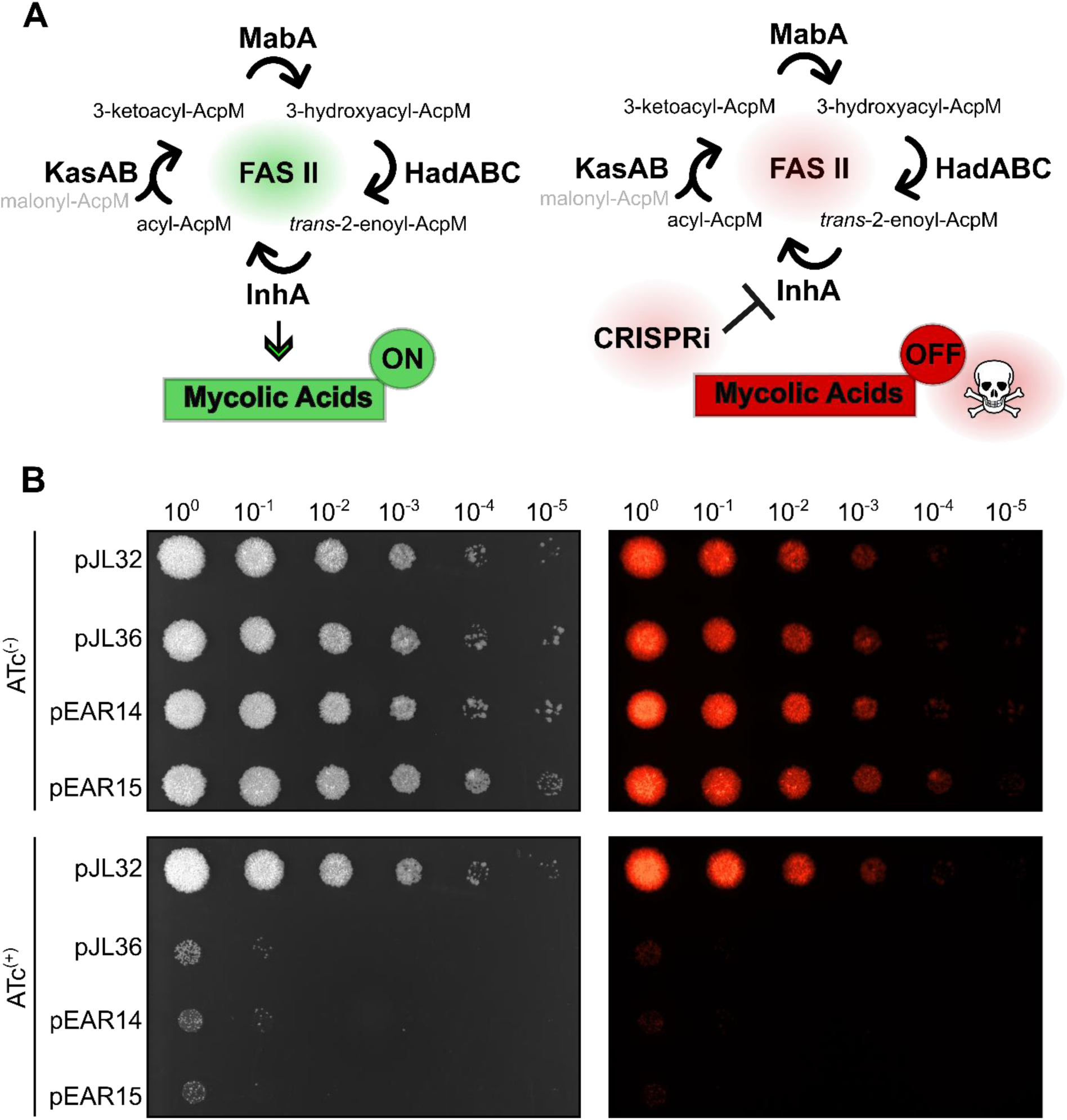
CRISPRi-targeting of *inhA* confirms its essentiality and role as key drug target in mycobacteria. **A.** Schematic representation of the mycobacterial FAS-II system and contribution of InhA in mycolic acid biosynthesis. InhA plays a critical role by fuelling the pool of mature meromycolic acid required for cell-wall biosynthesis and maintenance. Accordingly, CRISPRi-mediated genetic repression of *inhA* should result in mycobacterial growth arrest. **B.** CRISPRi-targeting of *inhA* expression in *Msmeg* confirms its essentiality *in vitro*. Serial dilutions of exponentially growing *Msmeg* recombinant strains pJL32, pJL36, pEAR14 and pEAR15 ranging from 10^6^ CFU to 10^1^ CFU were spotted onto 7H10 agar media either in the absence (top panel) or presence (bottom panel) of 100 ng. mL^-1^ of ATc. Bright light (left) and their corresponding red fluorescent profiles (right) are shown.

After transformation and selection of recombinant clones, validation of CRISPRi-mediated *inhA* silencing was performed using a well-established spot dilution assay on 7H10 solid media in the absence or presence of inducer, 100 ng·mL^−1^ of anhydrotetracycline (ATc) [34]. The assay included two additional control strains. A non-targeting control harboring the empty vector pJL32, which encodes solely the sgRNA scaffold without a gene-specific targeting sequence and a positive control of repression pJL36, which carries an integrative CRISPRi vector that successfully targets the expression of *mmpL3*, which encodes the essential trehalose monomycolate translocase [35–38]. Control condition in the absence of ATc revealed that all the strains grew as expected with no detectable defect (Fig. 1B). In addition, in the presence of ATc, no growth defect was observed for the non-targeting strain pJL32 confirming that the presence of ATc, dCas9 and sgRNA-scaffold alone expression did not impair growth (Fig. 1B). In contrast, analysis of pEAR14 and pEAR15 phenotypes, showed a significant 4 to 5-log_10_ decrease in bacterial growth, similar to the one obtained for the control strain pJL36, showing that both CRISPRi vectors targeting *inhA* are functional with efficient repression leading to mycobacterial growth inhibition (Fig. 1B).

### Genetic engineering of conditional and reversible *Msmeg* hypomorphs resistant to InhA-targeting compounds

In order to develop an inducible and reversible genetic system that triggers INH resistance, we capitalized on our CRISPRi-based tools to genetically engineered three distinct strains in which we repress *katG* (*MSMEG*_6384), the major catalase-peroxidase required for INH activation [39] (Fig. 2A). Accordingly, recombinant strains pWA2, pWA5 and pWA8, were designed and generated to silence *katG* with different sgRNA sequences (Fig. S1). Then, INH susceptibility of the three strains was assessed on 7H10 agar plates containing increasing concentrations of INH ranging from 0 to 64 µg·mL⁻¹ in the absence or presence of 100 ng·mL^−1^ of ATc and further compared to the pJL32 reference strain (Fig. 2B).

**Figure 2:**
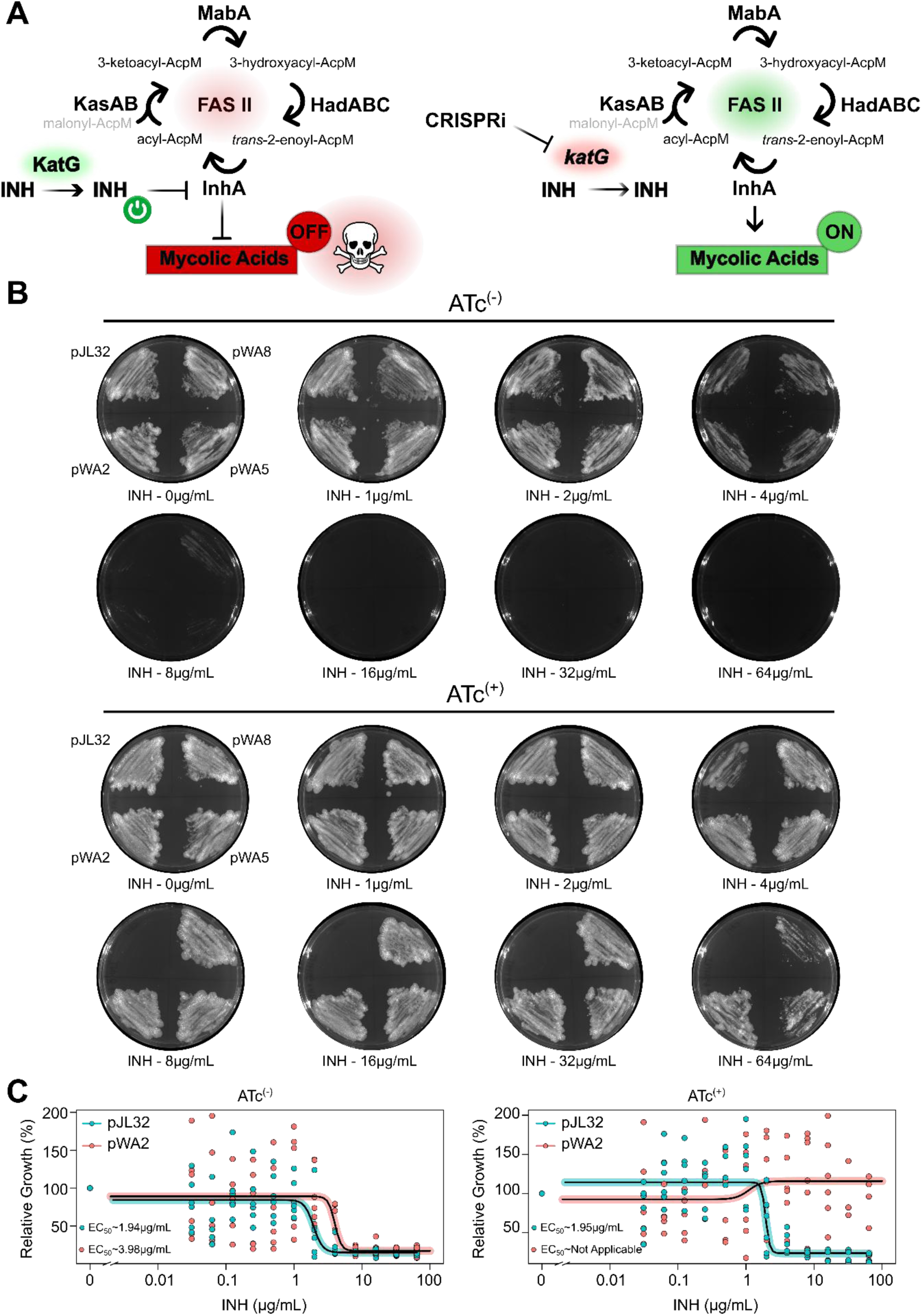
CRISPRi-targeting of *katG* results in conditional resistance to the front-line drug INH. **A.** Schematic representation of *katG* repression within the mycobacterial FAS-II system and potential effect on the level of mature meromycolic acid required for cell-wall biosynthesis and maintenance. Accordingly, CRISPRi-mediated genetic repression of *katG* should impair the bioactivation of INH, in its active form and therefore suppress its antibacterial activity. **B.** CRISPRi-targeting of *katG* expression in *Msmeg* causes INH resistance *in vitro* when assessed on agar plates. Approximately, 10^6^ CFU of exponentially growing *Msmeg* recombinant strains pJL32, pWA2, pWA5 and pWA8, were spotted and further evenly spread with a sterile loop on their corresponding quadrant of 7H10 agar plates containing 50 µg·mL⁻¹ of kanamycin, and increasing concentrations (ranging from 0 to 64 µg·mL⁻¹) of INH. Plates were done in the absence of inducer (ATc^(−)^ - top panel) or in the presence of 100 ng·mL⁻¹ of inducer (ATc^(+)^ - bottom panel). For clarity, only bright light acquisitions of the plates are displayed. Results are representative of two biological replicates. **C.** CRISPRi-targeting of *katG* expression in *Msmeg* causes INH resistance *in vitro* when assessed in broth. Comparative analysis of INH-mediated growth inhibition against *Msmeg* pJL32 (cyan) & pWA2 (pink) in Middlebrook 7H9 medium in the absence (ATc^(−)^ - left panel) or presence of 100 ng·mL⁻¹ of inducer (ATc^(+)^ - right panel). Dose-response analysis displayed were obtained following a 4-parameter nonlinear logistic regression and EC_50_ were determined accordingly. Results are representative of two biological replicates.

In the absence of ATc, the growth of pJL32, pWA2, pWA5 and pWA8 was comparable, with an observable complete growth inhibition achieved at 16 µg·mL⁻¹ of INH. In the presence 100 ng·mL^−1^ of ATc, pJL32 growth profile remained identical to its uninduced control condition. However, strains pWA2, pWA5 and pWA8 showed a substantial increase in INH resistance by still displaying very high-density growth at INH concentrations of up to 64 µg·mL⁻¹ (Fig. 2B). These results confirmed the successful repression of *katG* which confers high-level resistance towards INH. The pWA2 strain, which exhibited the strongest phenotype, was therefore selected for further validation in standard antibiotic susceptibility assays.

Hence, we confirmed CRISPRI-mediated alteration of antibiotic efficacy in 7H9 broth using the 96-well plate microdilution approach. Accordingly, bacterial growth was monitored at optical density 600nm (OD_600nm_) across increasing concentrations of INH from 0 to 64 µg·mL⁻¹. In the absence of ATc, both the control strain pJL32 and the *katG*-targeting strain pWA2 exhibited similar growth inhibition profiles, with INH half maximal effective concentration (EC_50_) of 1.94 and 3.98 µg·mL⁻¹ respectively and an observed MIC ranging between 4 and 8 µg·mL⁻¹ across replicates (Fig. 2C).

In the presence of ATc, pJL32 remained susceptible to INH with EC_50_ determined at 1.95 µg·mL⁻¹ and an MIC observed around 8 µg·mL⁻¹. In contrast, the pWA2 *katG-*hypomorph, showed a strong resistance phenotype with very high OD_600_ readings across the entire range of INH concentrations tested, confirming an ATc inducible resistant-phenotype to INH (Fig. 2C).

We then applied an identical strategy to target *ethA* and therefore modulate ETH susceptibility (Fig. 3A). Hence, we engineered three CRISPRi plasmids with distinct sgRNA-coding sequences to repress *ethA* expression (Fig. S1). The corresponding pEAR9, pEAR10 and pEAR11 recombinant strains were generated, and antibiotic susceptibility profiling was assessed on 7H10 agar plates using increasing concentrations of ETH ranging from 0 to 64 µg·mL⁻¹ in the absence or presence of 100 ng·mL^−1^ of ATc and further compared to the pJL32 reference strain (Fig. 3B).

**Figure 3:**
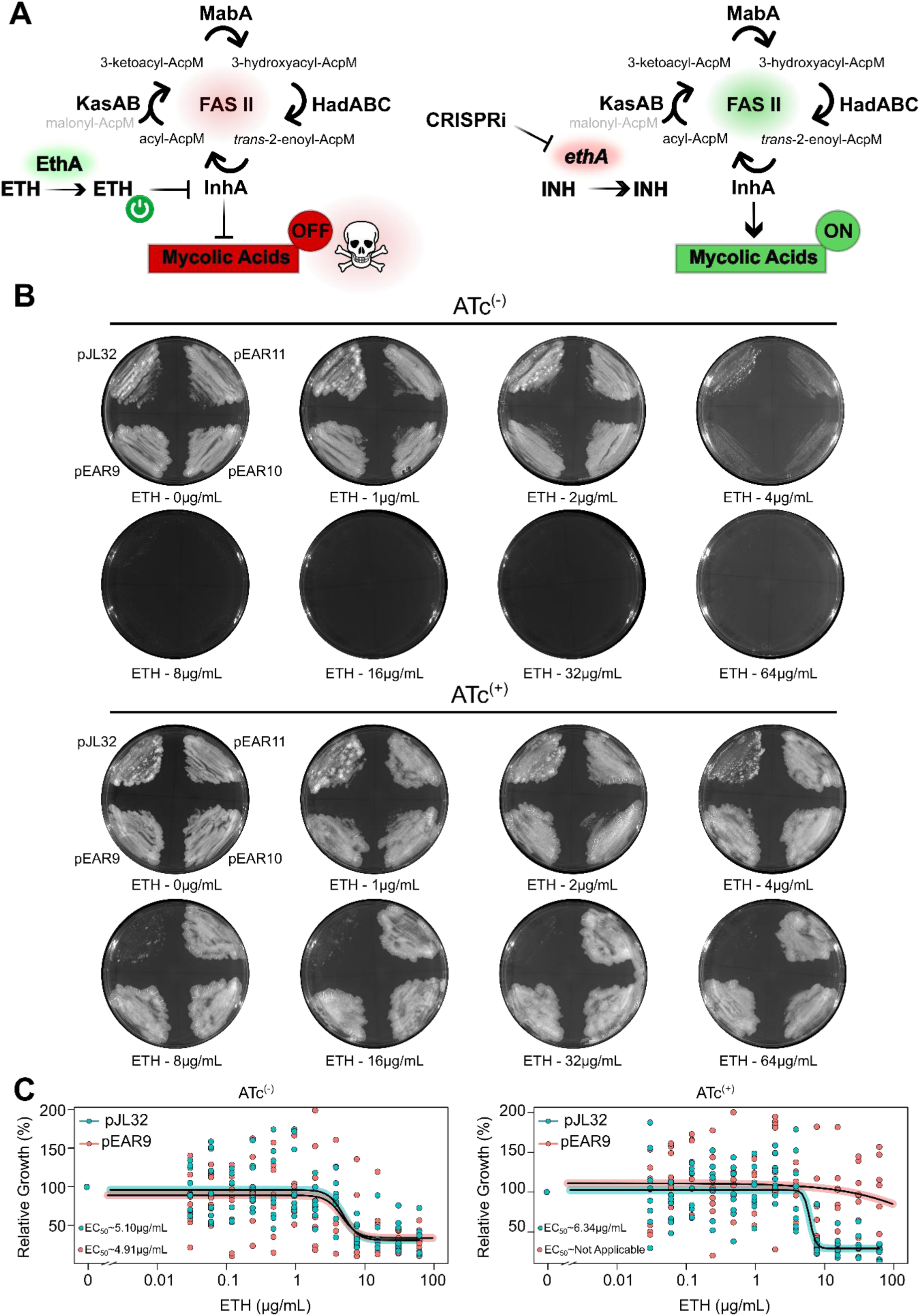
CRISPRi-targeting of *ethA* causes conditional resistance to ETH. **A.** Schematic representation of *ethA* repression within the mycobacterial FAS-II system and potential effect on the level of mature meromycolic acid required for cell-wall biosynthesis and maintenance. Accordingly, CRISPRi-mediated genetic repression of *ethA* should impair the bioactivation of ETH in its active form and therefore suppress its antibacterial activity. **B.** CRISPRi-targeting of *ethA* expression in *Msmeg* causes ETH resistance *in vitro* when assessed on agar plates. Approximately, 10^6^ CFU of exponentially growing *Msmeg* recombinant strains pJL32, pEAR9, pEAR10 and pEAR11, were spotted and further evenly spread with a sterile loop on their corresponding quadrant of 7H10 agar plates containing 50 µg·mL⁻¹ of kanamycin, and increasing concentrations (ranging from 0 to 64 µg·mL⁻¹) of ETH. Plates were done in the absence of inducer (ATc^(−)^ - top panel) or in the presence of 100 ng·mL⁻¹ of inducer (ATc^(+)^ - bottom panel). For clarity, only bright light acquisitions of the plates are displayed. Results are representative of two biological replicates. **C.** CRISPRi-targeting of *ethA* expression in *Msmeg* causes ETH resistance *in vitro* when assessed in broth. Comparative analysis of ETH-mediated growth inhibition against *Msmeg* pJL32 (cyan) & pEAR9 (pink) in Middlebrook 7H9 medium in the absence (ATc^(−)^ - left panel) or presence of 100 ng·mL⁻¹ of inducer (ATc^(+)^ - right panel). Dose-response analysis displayed were obtained following a 4-parameter nonlinear logistic regression and EC_50_ were determined accordingly. Results are representative of two biological replicates.

In the absence of ATc, the growth inhibition baseline for the non-targeting pJL32 strain sat around 16 to 32 µg·mL⁻¹ of ETH. Under the same conditions, *ethA*-silenced strains showed a similar susceptibility profile, with no detectable growth above 16-32 µg·mL⁻¹ (Fig. 3B). When assessed on ATc-containing media, pJL32 remained inhibited by ETH, around 16-32 µg·mL⁻¹, while the three *ethA*-repressed strains showed hyper-resistance profiles with high-bacterial load even at 64 µg·mL⁻¹, the highest concentration tested. Comparison between the three hypomorphs revealed similar resistance profiles and pEAR9 selected and tested in broth to confirm this result. Thus, in 7H9 broth devoid of ATc, pJL32 and pEAR9 had similar growth inhibition profiles, with estimated EC_50_ of 5.10 and 4.91 µg·mL⁻¹ respectively. However, in the presence of ATc, dose-response experiments showed that pJL32 conserved a similar susceptibility profile towards ETH with an estimated EC_50_ of 6.34 µg·mL⁻¹, while the *ethA*-silenced strain pEAR9 showed an ETH-resistant phenotype by maintaining bacterial growth across all ETH concentrations tested up to 64 µg·mL⁻¹.

In order to check that CRISPRi-mediated silencing of *katG* or *ethA* was specific towards INH and ETH/PTH respectively, we performed some cross resistance assays on solid medium (Fig. S2 & S3). As expected, *katG*-silencing (pWA2) had no effect on ETH/PTH susceptibility while compared with pJL32 strain in the absence or presence of ATc (Fig. S2). Likewise, *ethA*-silencing (pEAR9) had no effect on INH susceptibility, confirming the high-degree of resistance specificity of the generated hypomorphs (Fig. S3).

Altogether, these results confirm that CRISPRi-mediated repression enables the generation of ATc inducible resistant-phenotype to clinically relevant drugs with a great specificity while targeting pro-drug activation pathways.

### Fine tuning of prodrug activation through genetic engineering impact antibiotic vulnerabilities

With the recent development of chemical boosters that enhance prodrug bioactivation *via* binding inhibition of transcriptional repressors [18, 19], we hypothesized that inducible genetic repression of transcriptional repressors using CRISPRi should have effects very close to the ones observed with chemical boosters. As such, we postulated that CRISPRI-mediated repression of *ethR*, the transcriptional repressor that controls *ethA*, should boost *ethA* transcription, therefore leading to a greater level of EthA, that in turn will increase the bioactivation of ETH and lead to an increased susceptibility (Figure 4A).

**Figure 4:**
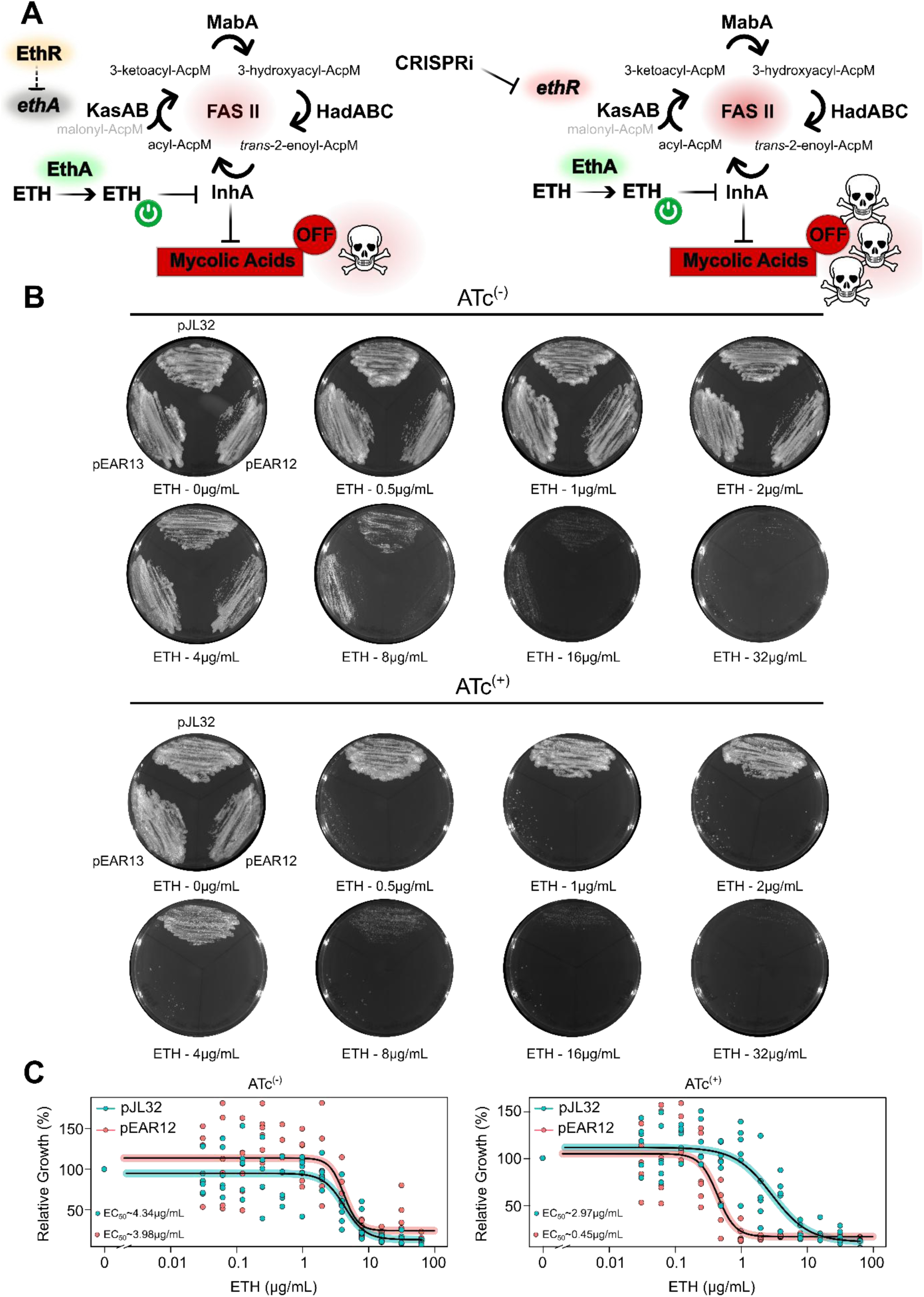
CRISPRi-targeting of *ethR* results in conditional hypervulnerability to ETH. **A.** Schematic representation of *ethR* repression within the mycobacterial FAS-II system and potential effect on the level of mature meromycolic acid required for cell-wall biosynthesis and maintenance. Accordingly, CRISPRi-mediated genetic repression of *ethR* should impair *ethA* repression, increase the bioactivation of ETH in its active form and therefore result in a more potent antibacterial activity. **B.** CRISPRi-targeting of *ethR* expression in *Msmeg* causes ETH hypervulnerability *in vitro* when assessed on agar plates. Approximately, 10^6^ CFU of exponentially growing *Msmeg* recombinant strains pJL32, pEAR12 and pEAR13, were spotted and further evenly spread with a sterile loop on their corresponding quadrant of 7H10 agar plates containing 50 µg·mL⁻¹ of kanamycin, and increasing concentrations (ranging from 0 to 64 µg·mL⁻¹) of ETH. Plates were done in the absence of inducer (ATc^(−)^ - top panel) or in the presence of 100 ng·mL⁻¹ of inducer (ATc^(+)^ - bottom panel). For clarity, only bright light acquisitions of the plates are displayed. Results are representative of two biological replicates. **C.** CRISPRi-targeting of *ethR* expression in *Msmeg* causes ETH hypervulnerability *in vitro* when assessed in broth. Comparative analysis of ETH-mediated growth inhibition against *Msmeg* pJL32 (cyan) & pEAR12 (pink) in Middlebrook 7H9 medium in the absence (ATc^(−)^ - left panel) or presence of 100 ng·mL⁻¹ of inducer (ATc^(+)^ - right panel). Dose-response analysis displayed were obtained following a 4-parameter nonlinear logistic regression and EC_50_ were determined accordingly. Results are representative of two biological replicates.

To test this, we generated two CRISPRi constructs, pEAR12 and pEAR13, each expressing distinct sgRNA-coding sequences targeting *ethR* (Fig. 4B; Fig. S1). The susceptibility to ETH for control strain pJL32 and *ethR*-silenced strains pEAR12 and pEAR13 was tested on 7H10 solid media supplemented with ETH at 0, 0.5, 1, 2, 4, 8, 16, and 32 µg·mL⁻¹. In the absence of the ATc, pJL32 showed a basal level of susceptibility to ETH, with strong growth inhibition at 16 µg·mL⁻¹and some small residual colonies still visible at 32 µg·mL⁻¹. Without induction, both pEAR12 and pEAR13 exhibited phenotypes indistinguishable from the pJL32 control, indicating that CRISPRi repression was not active under uninduced conditions. Of note, these inhibition results are in perfect accordance with results obtained previously using pJL32 and pEAR9, pEAR10 and pEAR11 with MIC approximating 16-32 µg·mL⁻¹ on solid media (Fig. 3B).

Upon ATc addition, the susceptibility profile of the pJL32 control strain remained unchanged. In contrast, both *ethR*-silenced strains showed hypersensitivity to ETH, where complete growth inhibition was evident at ETH concentrations as low as 0.5-1 µg·mL⁻¹, with pEAR12 being the most potent construct that displayed the strongest vulnerability. To quantitatively validate these findings, susceptibility assays were repeated in 7H9 broth with ETH concentrations ranging from 0 to 64 µg·mL⁻¹ by comparing the mutant strain pEAR12 to the pJL32 control strain (Fig. 4C). Without ATc, pJL32 and pEAR12 exhibited nearly identical growth inhibition profiles, with estimated ETH EC_50_ of 4.34 and 3.98 µg·mL⁻¹, respectively. Upon ATc induction, pJL32 control strain still displayed a comparable inhibition growth profile as its uninduced control conditions with an EC_50_ of 2.97 µg·mL⁻¹, while pEAR12 showed a drastic hyper vulnerability with an EC_50_ of 0.45 µg·mL⁻¹, resulting in an 8.8-fold reduction while comparing pEAR12 EC_50_ values in the presence and absence of ATc (Fig. 4C).

Finally, to complement this approach, we also generated pJL32-derived recombinant strains that harbor hygromycin-resistant episomal vectors, compatible with our CRISPRi-constructs. Hence, we cloned *ethA*, *ethR* or the control *mWasabi* under the strong constitutive promoter Psmyc, in order to constitutively express these genes and further determine their impact on ETH susceptibility (Fig. S4). Analysis of the susceptibility profiles of these recombinant strains on 7H10 agar plates, showed that *ethA* overexpression poorly affect susceptibility in comparison to the mWasabi-expressing control strains (Fig. S4). This suggests that basal transcriptional levels of *ethA*, controlled via EthR, are likely sufficient to create a vulnerability towards ETH in *Msmeg*, and that increasing production of EthA through Psmyc-driven expression minimally affects antibiotic efficacy. On the other hand, constitutive overexpression of *ethR* increasing antibiotic resistance on 7H10 media and in 7H9 broth by 3-fold as displayed in Figure S3.

All these experiments were further expanded to *Mtb* using the pJL34 backbone to further validate this approach (Fig. S1). Hence, we generated *Mtb* hypomorphs that display both hyper-resistant and hyper-susceptible profiles towards InhA-targeting compounds, by knocking-down *katG* (Fig. S5), *ethA* (Fig. S6) and *ethR* (Fig. 5) respectively, while compared to the control recombinant harboring the pJL34 empty vector. Data displayed as Fig. S5 & Fig. S6 showed that repression of *katG* or *ethA* increased the resistance towards INH and ETH on ATc-containing medium only. Thus, confirming that such inducible tools can be used to mimic clinical resistance in the tubercle bacilli. On the contrary, targeted repression of *ethR* led to a drastic decrease in susceptibility which was clearly observable on agar plates upon ATc induction (Fig. 5). This conditional, hyper-vulnerable phenotype to ETH matched the one obtained in *Msmeg* in Fig. 4 and was further studied in broth. Antibiotic susceptibility profiling of the *Mtb* pEAR21 strain was performed in the presence of increasing concentration of ETH, without (Fig. 5A – top panel) or with inducer (Fig. 5A – bottom panel). Determination of EC_50_ values through dose-response analysis and log-logistic regression demonstrated that induction of the CRISPRI system led to a substantial 10-fold decrease from 1.29 µg·mL⁻¹ to 0.12 µg·mL⁻¹ confirming this EthR-dependent hyper-vulnerable phenotype.

**Figure 5:**
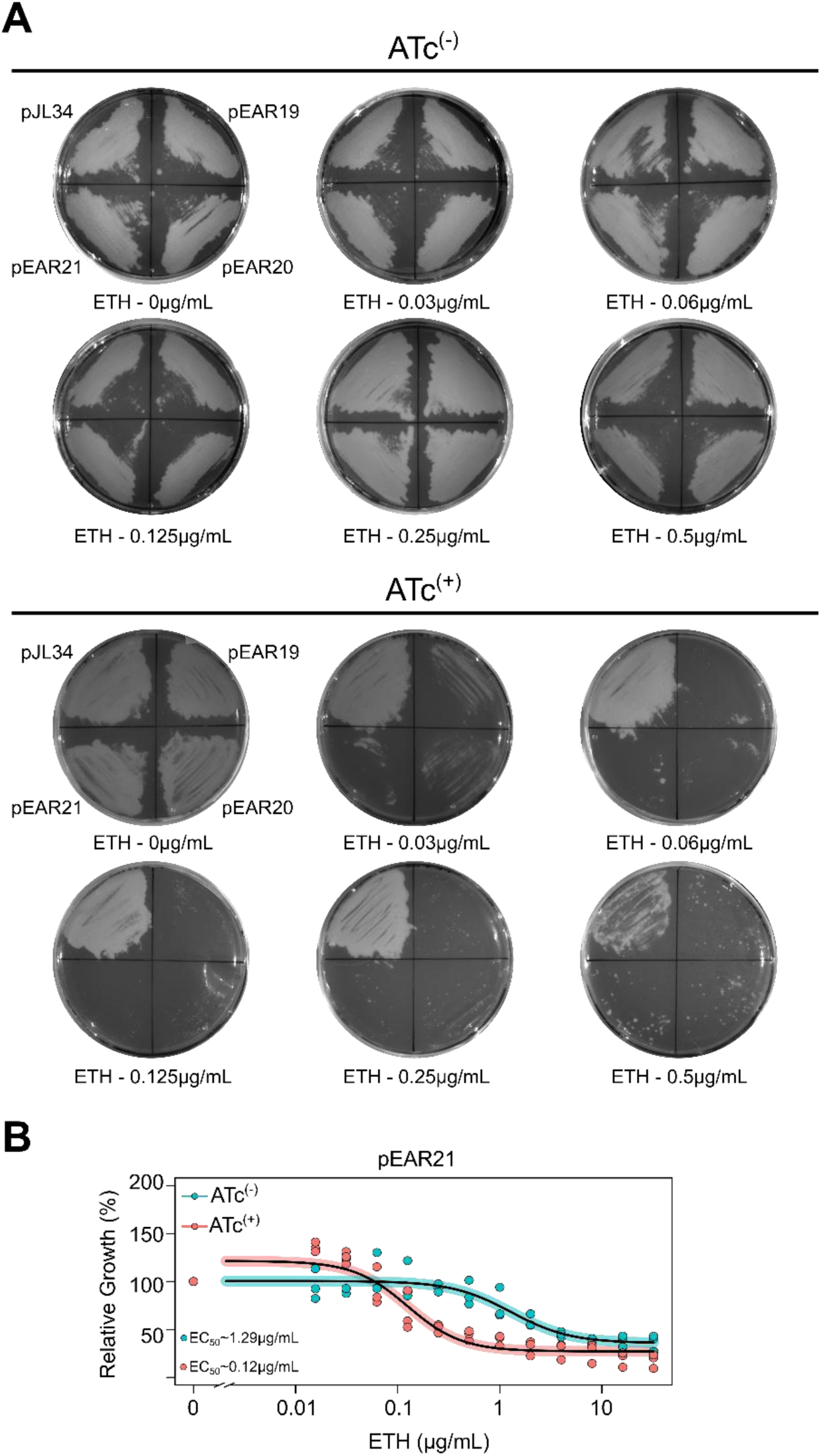
CRISPRi-based repression of *ethR* is also functional in *Mtb* and results in conditional hypervulnerability to ETH. **A.** CRISPRi-targeting of *ethR* expression in *Mtb* causes ETH hypervulnerability *in vitro* when assessed on agar plates. Approximately, 10^5^ CFU of exponentially growing *Mtb* recombinant strains pJL34, pEAR19, pEAR20 and pEAR21 were spotted and further evenly spread with a sterile loop on their corresponding quadrant of 7H10 agar plates containing 10% OADC enrichment, 50 µg·mL⁻¹ of kanamycin, and increasing concentrations (ranging from 0 to 0.5 µg·mL⁻¹) of ETH. Plates were done in the absence of inducer (ATc^(−)^ - top panel) or in the presence of 100 ng·mL⁻¹ of inducer (ATc^(+)^ - bottom panel). For clarity, only bright light acquisitions of the plates are displayed. Results are representative of two biological replicates. **B.** CRISPRi-targeting of *ethR* expression in *Mtb* causes ETH hypervulnerability *in vitro* when assessed in broth. Comparative analysis of ETH-mediated growth inhibition against *Mtb* pEAR21 in complete Middlebrook 7H9 medium in the absence (ATc^(−)^ - cyan) or presence of 100 ng·mL⁻¹ of inducer (ATc^(+)^ - red). Dose-response analysis displayed were obtained following a 4-parameter nonlinear logistic regression and EC_50_ were determined accordingly. Results are representative of three biological replicates.

These results in *Mtb* confirm that our genetic platform provides a reliable system in mycobacteria that enables to chemically induce fine-tuning of antibiotic efficacy by modulating vulnerabilities towards resistance or susceptibility. More importantly, these results confirm that this genetic-based modulation of antibiotic efficacy by overexpressing or repressing specific target genes could have a wide range of application for our community.

## Discussion

As the global TB pandemic continues to be a major health issue, the high number of RR-TB and MDR-TB cases underscores the urgent need for novel strategies to eradicate TB, including the development of new drug regimens [1, 2]. While several progresses have been made and new regimens are giving extremely promising results, we still need to further expand our arsenal of drugs, especially if we aim at achieving the goals set by the End TB Strategy for 2035 [1, 2].

To find new molecules that are active against drug-resistant strains through tomorrow’s drug discovery screening, it is crucial to develop new approaches and genetic tools that facilitate the next generation of whole-cell phenotypic screening assays. Conventional genetic methods, such as isolation of loss-of-function antibiotic resistant mutants or the generation of knockout mutants are extremely valuable [19], but their development and validation can be extremely time consuming. Moreover, such tools lack the temporal control and reversibility that might be required to study dynamic biological processes.

In this work, we demonstrate that CRISPRi overcomes some of these limitations, and report a fast framework to generate recombinant strains that enables an inducible, tunable, and reversible targeted-gene silencing in mycobacteria. Our results align with previous studies highlighting CRISPRi as a powerful tool for gene-silencing and functional genomics in bacteria, including *Mtb* [31, 32, 40].

Accordingly, we first targeted the well-characterized therapeutic target InhA, which is essential for mycobacterial growth, as a proof of concept to validate the functionality of our CRISPRi approach [31, 41]. Indeed, the essentiality of *inhA* was assessed through CRISPRi-mediated silencing, which resulted in a 4–5 log_10_ reduction in bacterial growth, consistent with its critical role in mycolic acid biosynthesis [4, 7]. This is in perfect agreement with our recent data showing that *mmpL3* or *rpoB* could be also efficiently targeted by our CRISPRi system [33, 34]. Therefore, it acts as a very reliable method for studying essential genes in mycobacteria, which are often recalcitrant to traditional genetic manipulation.

Clinical resistance to INH and ETH/PTH in *Mtb* is primarily driven by loss-of-function mutations in *katG* and *ethA*, respectively [6, 11, 13, 17]. Unsurprisingly, CRISPRi-mediated repression of *katG* and *ethA* in *Msmeg* phenocopied clinical resistance, with strains exhibiting high-level resistance up to 64 µg·mL⁻¹ only upon induction with ATc. These findings are consistent with the well-established prodrug activation paradigm, wherein KatG and EthA are required to convert INH and ETH into their active forms [6, 11, 13, 17]. Moreover, our results corroborate the findings from Ando *et al*. who demonstrated that specific mutations in the *furA-katG* intergenic region decreased *katG* expression and conferred INH resistance in *Mtb* [42, 43]. Based on this observation, we could suggest that the tunable feature of CRISPRi, could act as a powerful tool to mimic some of these mutations that are often complicated to select *in vitro* or generate through genome-editing. Hence, targeting of KatG through distinct PAM or sgRNA sequences, with different level of strength and/or increasing concentration of inducers, could enable the generation of a diverse panel of recombinant strains spanning a broad range of resistance phenotypes, thereby providing experimental models that better reflect the spectrum of resistance observed in clinical isolates.

Work performed in this study also confirms the pioneering work from the Rock’s group; showing that targeted CRISPRi or whole-genome assays act as an extremely powerful tool to identify new targets and/or mode of action [44, 45]. Indeed, Gouzy and colleagues, have recently applied genome-wide CRISPRi profiling associated to comparative genomics and high-resolution metabolomics, to unravel a previously unknown resistance mechanism towards the prodrug PZA [45]. Indeed, this unique approach enabled to show that within acidic microenvironments mimicking host infection, α-ketoglutarate efflux *via* Rv2571c potentiates cytoplasmic acidification caused by PZA. Consequently, loss-of-function mutations in Rv2571c abrogate this effect, yielding resistance *in vitro*, *in vivo* and are positively selected in patient isolates. This highlights the strength of the approach, which allows, similarly to TnSeq, to identify hyper-vulnerable phenotype that could only be detected through whole-genome screening.

Another important result from our work resides in our cross-resistance assays (Fig. S2 & Fig. S3), which demonstrated that *katG* repression did not affect ETH/PTH susceptibility, and *ethA* repression did not alter INH susceptibility. This specificity confirms that our CRISPRi hypomorphs accurately model the mechanistic basis of drug resistance to INH and ETH/PTH in mycobacteria, without major off-target effects. These results also highlight the modularity of our approach, as CRISPRi can be tailored to target distinct bioactivation pathways independently. Accordingly, it would be interesting to generate a novel subset of CRISPRi constructs that contains two independent sgRNA that enable to simultaneously target both *katG* and *ethA*; resulting in a strain that is INH/ETH/PTH resistant through impairment of two distinct bioactivation pathways.

One of the most striking findings of this study is the bidirectional modulation of antibiotic susceptibility achieved through CRISPRi [44, 45]. While repression of *katG* or *ethA* conferred resistance, silencing of the transcriptional repressor *ethR* led to hyper-susceptibility to ETH, with an 8.8-fold reduction in EC_50_ (from 3.98 µg·mL⁻¹ to 0.45 µg·mL⁻¹) upon ATc induction. This phenomenon mirrors the action of chemical boosters like alpibectir, which inhibits EthR to derepress *ethA* expression, thereby enhancing ETH bioactivation [18, 21, 22]. Our genetic approach thus validates EthR as a complementary drug target and demonstrates that CRISPRi can emulate the effects of small-molecule inhibitors, offering a complementary strategy for drug development. The hypersusceptibility phenotype observed in *ethR*-silenced strains confirm that transcriptional derepression of *ethA* is sufficient to hyper-sensitize mycobacteria to ETH. Furthermore, we validated this approach and phenotype in *Mtb* demonstrating the translatability of our system to clinically relevant pathogens.

Overall, this study establishes a genetic platform for high-throughput screening of anti-TB compounds, particularly those targeting InhA or prodrug activation pathways. By generating conditional hypomorphs, we can easily identify novel InhA inhibitors that completely bypass the need for bioactivation by KatG or EthA. As such, this technology opens new avenues for identifying new drugs, new targets and/or mode of action, but might also facilitate preclinical testing of combination therapies that pair CRISPRi-mediated gene repression with existing antibiotics [44] within *in vitro* conditions that mimic the physiological context. Moreover, by identifying genetic or chemical perturbations that increase bacterial susceptibility to established anti-TB drugs, this approach could reveal cellular pathways and mechanisms that contribute to drug resistance, tolerance, and persistence, including vulnerabilities that may only become apparent under antibiotic pressure. Importantly, such sensitization targets could represent novel therapeutic opportunities, enabling the development of antibiotic potentiators or adjuvant therapies that enhance the activity of existing drugs rather than requiring the discovery of entirely new antibiotics. This strategy could therefore facilitate the rational identification of synergistic drug combinations, provide opportunities for drug repurposing, and potentially improve treatment efficacy against drug-tolerant or persistent bacterial populations.

## Conclusion

In summary, this work demonstrates that CRISPRi is a robust and versatile tool for modulating antibiotic susceptibility in mycobacteria by targeting genes involved in prodrug activation or coding transcriptional factors involved in their regulation. Building upon foundational work from the Rock laboratory, our findings and experimental toolkit establish a robust genetic framework for dissecting and overcoming resistance mechanisms that could be of broad interest. Indeed, by enabling targeted, inducible, and reversible gene silencing, CRISPRi offers a complementary approach to chemical boosters and a powerful platform for the next generation of anti-TB drug screenings.

## Material & Methods

### Bacterial strains and culture conditions

*M. smegmatis* mc^2^155 and *M. tuberculosis* ATCC 25177 H37Ra reference strains were used in this study. All mycobacterial strains were routinely grown in Middlebrook 7H9 broth (BD Difco, #271310) supplemented with 0.2% glycerol (Euromedex, #EU3550) and 0.05% Tween 80 (Sigma-Aldrich, #P1754). In the case of *M. tuberculosis*, 10% oleic acid, albumin, dextrose and catalase (OADC enrichment; BD Difco, #211886) was also added. All cultures were maintained at 37 °C with shaking.

All molecular biology experiments including plasmid selection, expansion and maintenance were performed in the reference strain *Escherichia coli* DH10B (Invitrogen). *E. coli* was grown into LB Broth (BD Difco; #244620) or in LB solid medium containing Bacteriological Agar (Euromedex; #1330). When required, hygromycin B (Toku-E, #H007) or kanamycin (Euromedex, #UK0010D) were used as selection markers for transformants on solid medium or for the culture maintenance of the recombinant strains. Final concentrations of 200 µg·mL⁻¹ and 50 µg·mL⁻¹were used respectively for *E. coli* while 50 µg·mL⁻¹ was used for *Msmeg* and *Mtb*.

### Plasmid constructions and generation of recombinant strains

All plasmids and primers used in this study are listed in supplementary information in Table S1 and Table S2.

Fluorescent-based CRISPRi plasmid backbones pJL32 (Addgene; #227429) and pJL34 (Addgene; #227431) were used for single-guide ribonucleic acid (sgRNA) insertion and gene knock-down [33]. Oligonucleotides targeting the non-template strand of the desired ORF to be targeted were designed using the Pebble database (https://pebble.rockefeller.edu/) following the recommendation of Bosch *et al*. [31]. H37Rv reference genome was used to design the constructs and further applied to its H37Ra counterpart. Accordingly, Rv annotation numbers were conserved for simplicity and the vectors developed can be used for both *Mtb* species. The CRISPRi backbones pJL32 and pJL34 were digested with BsmBI-v2 (NEB; #R0739L) and further purified using a Gel and PCR Clean-up kit (Macherey-Nagel; #740609.50).

To clone the sgRNA targeting sequences, complementary top and bottom oligonucleotides with appropriate sticky end overhangs, were resuspended, pooled and annealed following the protocol described by Wong *et al* [46]. Annealed oligos were dilute and ligated into the corresponding BsmBI-digested CRISPRi plasmid backbones using T4 DNA ligase (NEB; #M0202S). Chemically competent *E. coli* DH10B cells (Invitrogen) were transformed with 10–100 ng of ligation product and transformants were selected on LB agar supplemented with kanamycin (50 µg·mL⁻¹). Successful cloning was confirmed by Sanger sequencing (Eurofins Genomics) using the sequencing primer #P1. All primers (Integrated DNA Technologies, IDT) used to generate the sgRNA targeting sequences are listed in Table S1 (#P2–P33), and the targeting location of the sgRNA-dCas9 complex is schematized in Fig. S1.

To generate *Msmeg ethA* and *ethR* overexpression vectors, *ethA* and ethR were amplified by PCR from *Msmeg* mc^2^155 genomic DNA using #P34/#P35 or #P36/#P37 primers. Both PCR products were purified (NucleoSpin® Gel and PCR Clean-up, Macherey-Nagel) and further digested with PacI and HindIII restriction enzymes (New England Biolabs, #R0547L and #R3104L). Simultaneously, pUV15-pHGFP (Addgene plasmid #70045, kindly gifted by Sabine Ehrt) [47] was digested with PacI and HindIII to remove the pH-GFP coding sequence. Purified inserts and vector were then ligated using T4 DNA ligase to generate the pMSD45 and the pMSD46 vectors. Accordingly, pMSD45 and pMSD46 are hygromycin resistant episomal expression vectors harboring *ethA* and *ethR* genes respectively, cloned under the control of the Psmyc promoter.

All plasmids were transformed into electrocompetent *Msmeg* or *Mtb* cells as previously described by Goude *et al*. [48]. Briefly, a 500 mL culture of cells at optical density at 600 nm (OD_600_) 0.4-0.8 was harvested and washed five times in a sterile solution containing water, 10% glycerol and 0.05% Tween-80. The volume of washing solution was sequentially reduced (100, 50, 20, 10 mL), and the final pellet was resuspended in 5 mL. Then, 100 μL of electrocompetent cells were mixed with 500 ng to 1 μg of vector of interest in a pre-chilled 0.2 mm gap electroporation cuvette. *Msmeg* cells were subjected to a single pulse with 2.5 kV, 10 μF and pulse-controller resistance set at 600 Ω, while for *Mtb* resistance was set at 1000 Ω.

Subsequently, electroporated cells were resuspended in antibiotic-free complete 7H9 broth, and incubated at 37°C for 3–4 hours for *Msmeg* cells or 24 hours for *Mtb* cells with slow shaking (50 rpm). Transformants were selected on complete 7H10 agar supplemented with appropriate concentration of selection markers. For *Msmeg*, plates were incubated at 37°C for 4–7 days, while for *Mtb* they were incubated at 37°C for 21 days. When appliable, red/green fluorescent clones were selected using ChemiDoc^TM^ imaging system, inoculated into 5 mL of complete 7H9 broth containing the appropriate selection marker and grown at 37°C with shaking before being stored at −80°C in 20% glycerol.

### Analysis of CRISPRi-mediated silencing of *inhA* on solid media

Assessment of genetic essentiality was performed by performing serial dilutions of *Msmeg* recombinant strains and plated them on 7H10 media in the presence or the absence of the ATc inducer (Sigma-Aldrich, Saint-Quentin Fallavier, France; Supelco, Cat#37919) at final concentration of 100 ng·mL−1 as previously described by our group [33, 34]. Briefly, frozen *Msmeg* transformants stored at −80°C were thawed and grown for 48 hours in 7H9 broth with 50 µg·mL⁻¹ kanamycin, sub-cultured for 24 hours to reach exponential phase. Exponentially growing bacteria were normalized to an OD_600nm_ of 1 corresponding to approximately 1.10^8^ CFU per mL. Multiple 10-fold serial dilutions were performed, and 10 μL of each dilution (10^0^ to 10^−5^) was plated on each plate containing or not ATc. Plates were incubated at 37 °C, and visual inspection of growth was performed between days 2 and 4. Scanning of the plates was performed using the ChemiDoc™ MP Imaging System coupled with Image Lab software in two different settings: bright light acquisition (“White Epi Illumination” with “Standard Filter”), and red fluorescence acquisition (“Green Epi Illumination” setting with the “605/50 nm Filter”).

### Agar-based assessment of CRISPRi-mediated tuning of antibiotic susceptibility

Frozen *Msmeg* transformants stored at −80°C were thawed and grown for 48 hours in 7H9 broth with 50 µg·mL⁻¹ kanamycin, subcultured for 24 hours and normalized to OD_600_ of 1. The 7H10 agar plates containing kanamycin and increasing concentrations (0 to 64 µg·mL⁻¹) of either isoniazid (INH) (Sigma-Aldrich #I3377), ethionamide (ETH) (Sigma-Aldrich, #E6005) or prothionamide (PTH) (Sigma-Aldrich, #SMB00387) were prepared and virtually sectioned into two, three or four equal parts with each section assigned to a CRISPRI control or a CRISPRi hypomorph strain. Duplicates of the plates containing anhydrotetracycline (ATc) (Sigma-Aldrich, #37919) at 100 ng·mL⁻¹ were also poured and sectioned. A volume of 10 μL of a normalized culture of OD_600nm_ of 1, or alternatively 0.1, was spotted into its corresponding quadrant on the plate and evenly spread with a sterile loop. Inoculated plates in the presence or absence of ATc were then incubated at 37°C for 3-4 days.

A similar approach was used for *Mtb*, with slight modifications. Briefly, frozen *Mtb* transformants stored at −80°C were thawed and grown for 7 days in complete 7H9 broth containing 10% OADC enrichment and 50 µg·mL⁻¹ kanamycin. Saturated cultures were then sub-cultured for 5-7 days until reaching mid-exponential phase and further normalized to OD_600_ of 0.1. A volume of 10 μL of this normalized culture was spotted into its corresponding quadrant on 7H10 agar plates containing kanamycin, 10% OADC, and increasing concentrations of ETH or INH, and evenly spread with a sterile loop. Inoculated plates in the presence or absence of ATc were then incubated at 37°C for 15-21 days, before being imaged using the ChemiDoc^TM^ MP Imaging System coupled with Image Lab software as described above.

### Antibiotic susceptibility profiling assays in 7H9 broth using microdilution approach

Antibiotic susceptibility profiling of *Msmeg* and *Mtb* were performed using the 96-well microdilution approach as previously described [49, 50]. Briefly, all experiments were performed in standard Middlebrook 7H9 medium containing 0.2% glycerol and 0.05% Tween-80. For *Mtb,* 10% of OADC enrichment was added to the media. When required, kanamycin, hygromycin, or both selection markers were added to the media to maintain plasmid selection. Experiments were performed in the presence or absence of ATc to induce the CRISPRi system and assess targeted genetic repression on antibiotic efficacy.

Two-fold serial dilutions of the compound stock solutions were used to obtain a range of 12 concentrations in complete Middlebrook 7H9 broth. Exponential-phase mycobacterial cultures (OD_600_ of 1 ± 0.4) were adjusted to a bacterial density of 5 × 10^6^ CFU/mL in complete 7H9 medium. A volume of 100 μL of this inoculum was then dispensed into each well containing 100 μL of each serial dilution of the compound, achieving a final bacterial concentration of 2.5 × 10^6^ CFU/mL per well. The plate layout systematically included positive growth controls (i.e., inoculum without antibiotics) with and without ATc, vehicle growth controls (i.e. DMSO or ethanol), negative growth controls (i.e. using an additional control antibiotic to confirm general bacterial susceptibility), and sterility controls (i.e., medium only).

The 96-well flat-bottom Nunclon Delta Surface microplates with lid (Thermo-Fisher Scientific; #167008) were incubated at 37°C for 5 days for *Msmeg* and 14 days for *Mtb*. The minimum inhibitory concentration (MIC) of each antibiotic was visually determined in accordance with the EUCAST guidelines (https://www.eucast.org/mycobacteria/methodology-and-instructions/reference-method/). Results were confirmed by OD_600nm_ measurement using the TECAN spark 10M™ multimode microplate reader. Optical density measurements were used to determine a half-maximal effective concentration (EC_50_) using a four-parameter log-logistic nonlinear regression model (R Studio software, The R Project for Statistical Computing), as the concentration required to achieve 50% of the observed effect (i.e., mycobacterial growth inhibition). All the presented data have been obtained from experiments performed in technical triplicate at least on two or three distinct occasions.

## Acknowledgements

We would like to acknowledge all members of the Lipolysis and Bacterial Pathogenicity group and the LISM unit for continuous support and insightful discussions.

The work developed in our group is supported by the Centre National de la Recherche Scientifique (CNRS) and Aix-Marseille Université (AMU). PS received financial support from the CNRS Biologie, the Agence Nationale de Recherche sur le Sida et les Hépatites virales (ANRS/MIE) (project n°ANRS0358), the Agence Nationale de la Recherche (ANR) (ANR-24-CE15-2633 and Project ANR-25-ERCS-0022) and the French government under the France 2030 investment plan, as part of the Initiative d’Excellence d’Aix-Marseille Université - A*MIDEX and is part of the Institute of Microbiology, Bioenergies and Biotechnology - IM2B (AMX-19-IET-006). MSD PhD fellowship was funded by CONICET and her research stay was sponsored by the FEBS-IUBMB-PABMB PROBio-LatAm Fellowship Programme and the Compagny of Biologists Travelling Fellowship (#DMMTF25051802). JL PhD fellowship was funded by the Ministère de l’Enseignement Supérieur et de la Recherche. WA postdoctoral fellowship was funded by the foundation IHU Méditerranée Infection. PS has also received a FEBS Excellence Award 2023 to support this work and would like to personally thank the FEBS Excellence Awards and Fellowships Office for their continuous support. Special thanks are addressed to Alain Baulard for continuous support and his inspirational work on prodrug activation in mycobacteria.

The funders did not play a role in the study design, data collection and analysis, decision to publish, or preparation of the manuscript.

## Authors contribution

PS proposed, conceived and led the study. PS secured funding. GG and PS co-advised the PhD work of MSD. SC and PS co-advised the PhD work of JL. SC, CC and PS co-advised the work of WA. MM co-supervised the work of TB and VP with PS. ESR, MSD, MM, VP, TB, WA and JL performed the experimental work. ESR, MSD, MM, VP and PS edited the figures. All authors provided intellectual input by organizing, analysing and/or discussing data. PS and ESR, wrote the initial draft of the manuscript with input from all authors. All authors read the manuscript and provided critical feedback before its submission.

## Declaration of generative AI and AI-assisted technologies in the manuscript preparation process

During the preparation of this work the authors used https://chat.mistral.ai/ in order to perform some spell check, identify grammatical errors and improve the style and tone of the manuscript. After using this tool, the authors reviewed and edited the content as needed and take full responsibility for the content of this article.

## Data accessibility

Original plasmid vector pJL32 and pJL34 are available at (https://www.addgene.org/). Any additional vectors, strains or data that support the findings of this study are available upon reasonable request from the corresponding author at.

## Abbreviations

TB: Tuberculosis
*Mtb*: *Mycobacterium tuberculosis*
INH: isoniazid
ETH: ethionamide
PTH: prothionamide
CRISPRi: CRISPR interference
*Msmeg*: *Mycobacterium smegmatis*
RIF: rifampicin
EMB: ethambutol
PZA: pyrazinamide
RR-TB: rifampicin-resistant TB
MDR-TB: multidrug-resistant TB
ATc: anhydrotetracycline
OD_600nm_: Optical density at 600_nm_
EC_50_: half maximal effective concentration
sgRNA: single-guide ribonucleic acid
MIC: minimal inhibitory concentration

## Supporting Information

**Table S1.**
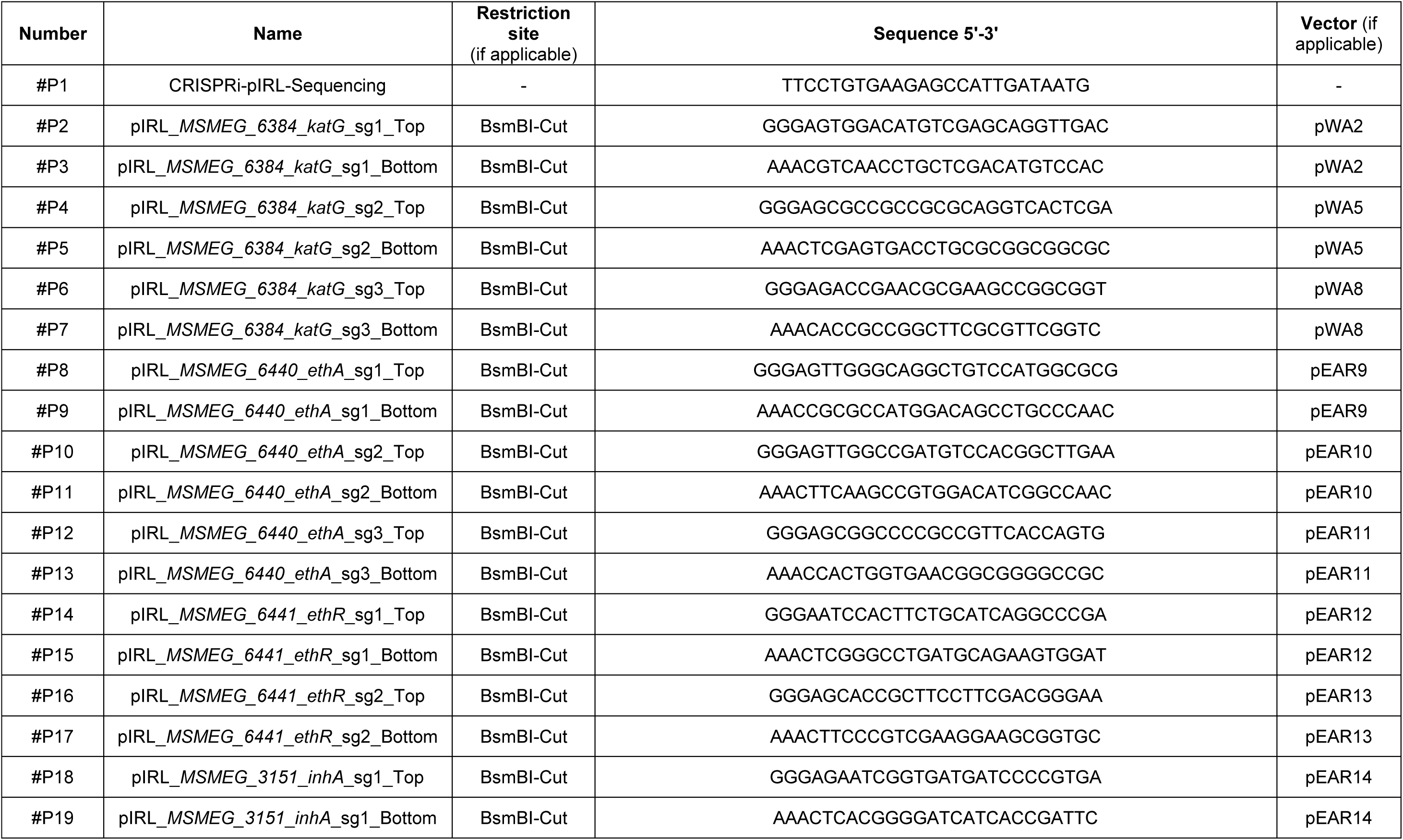
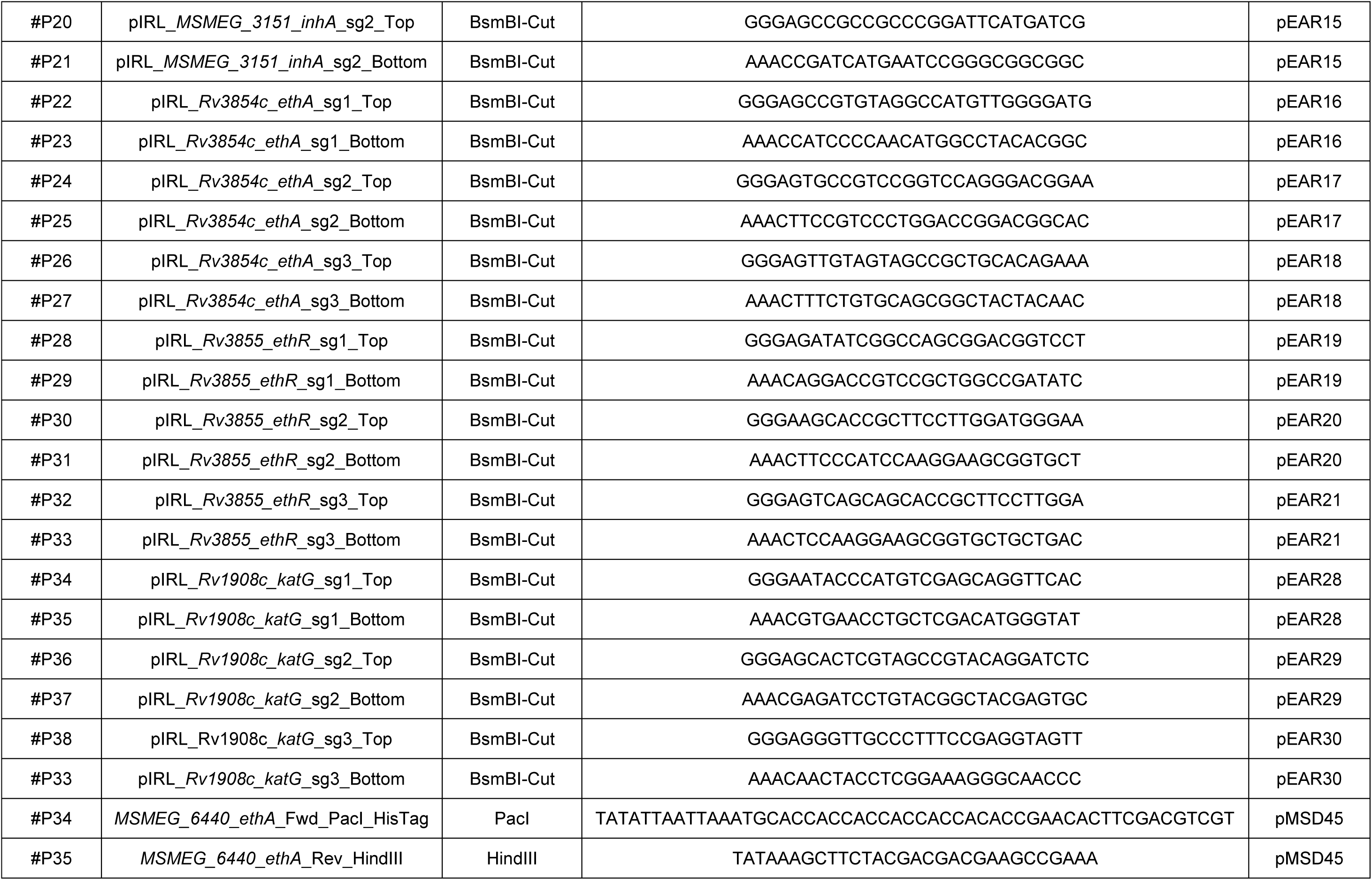
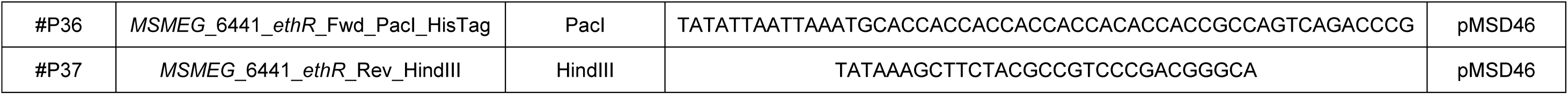
List of the primers used in this study to construct *M. smegmatis* & *M. tuberculosis* recombinant strains.

**Table S2.**
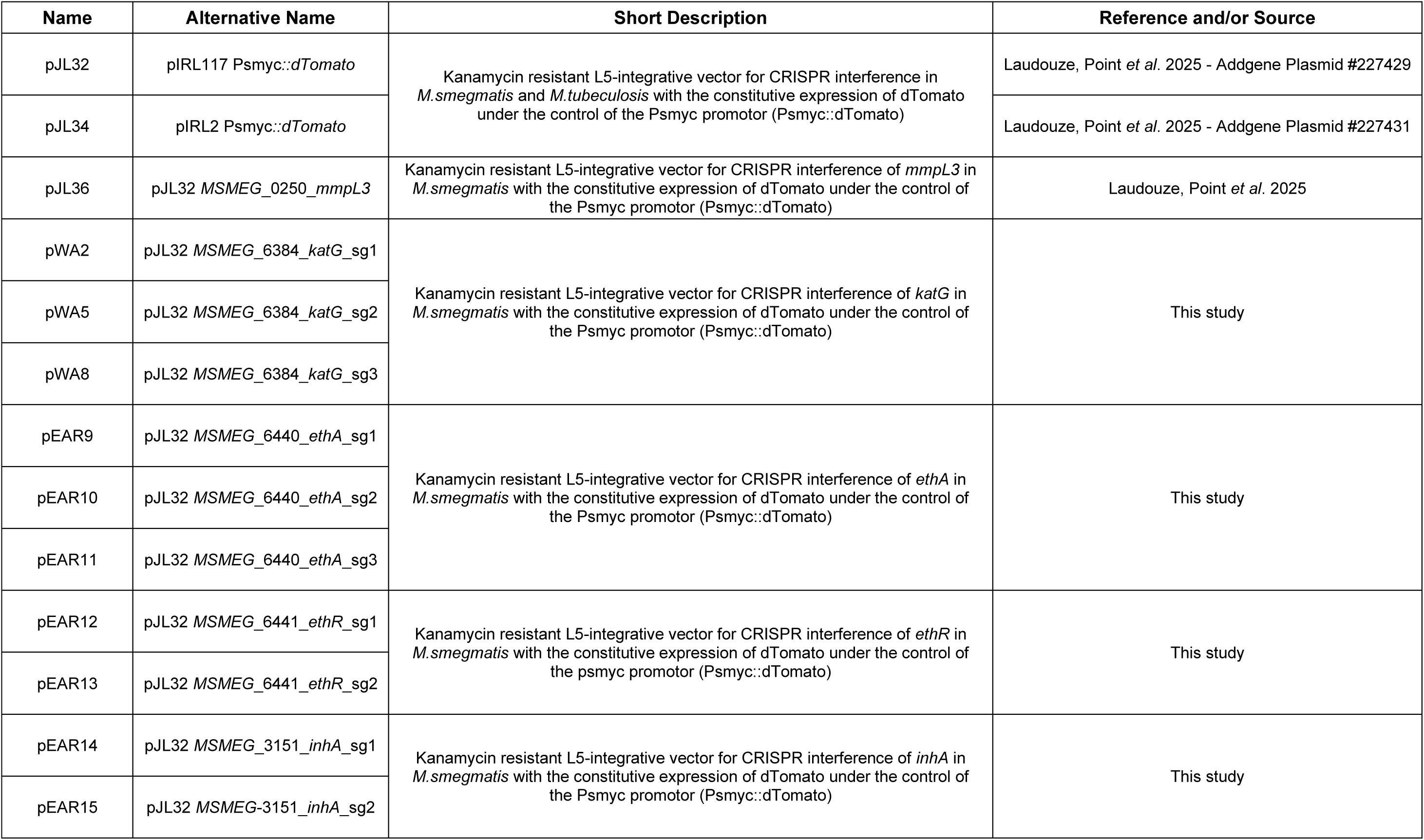
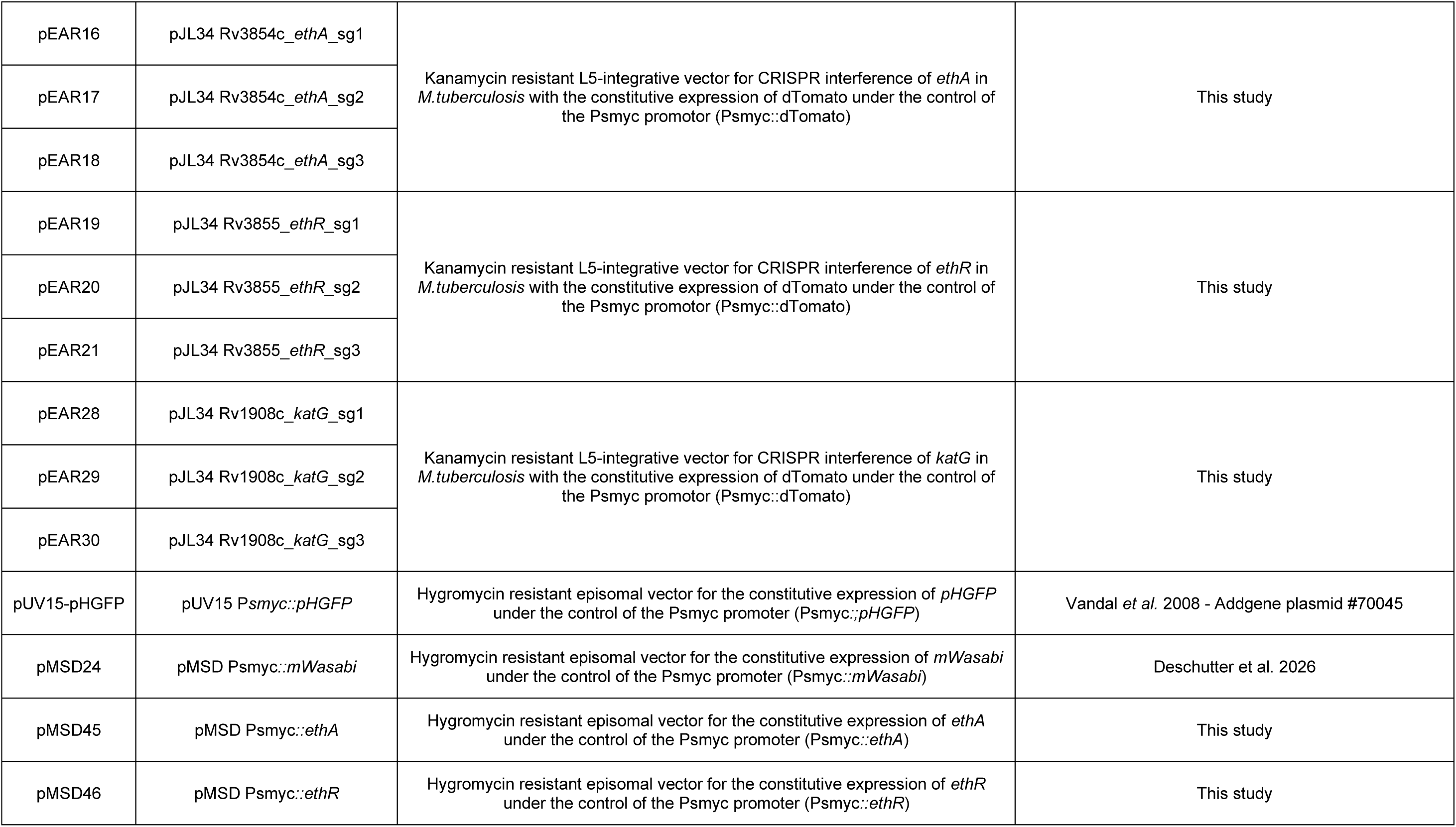
List of the plasmids used in this study.

**Figure S1.**
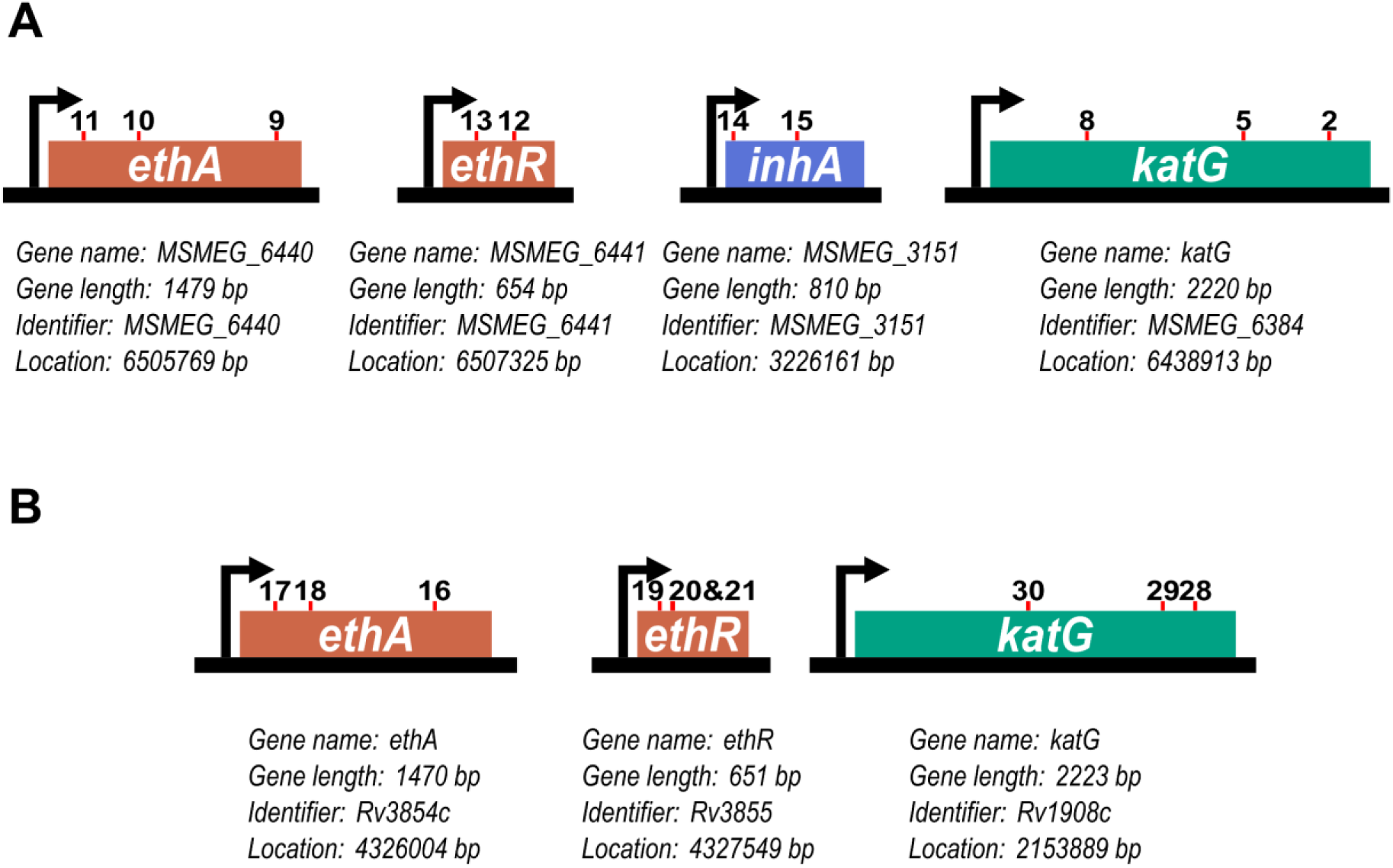
Schematic representation of the genes targeted via CRISPRi within this study and the respective positioning of the sgRNA-dCas9 complexes to perform silencing. **A.** Schematic representation of the positioning of the sgRNA-dCas9 complexes targeting *ethA*, *ethR*, *inhA* and *katG* in *Msmeg* mc^2^155 reference strain. Information about each gene and identifier are available under each scheme respectively. Information was obtained from the Mycobrowser database. **B.** Schematic representation of the positioning of the sgRNA-dCas9 complexes targeting *ethA*, *ethR*, and *katG* in *Mtb* H37Rv reference strain. Information about each gene and identifier are available under each scheme respectively. Information was obtained from the Mycobrowser database.

**Figure S2.**
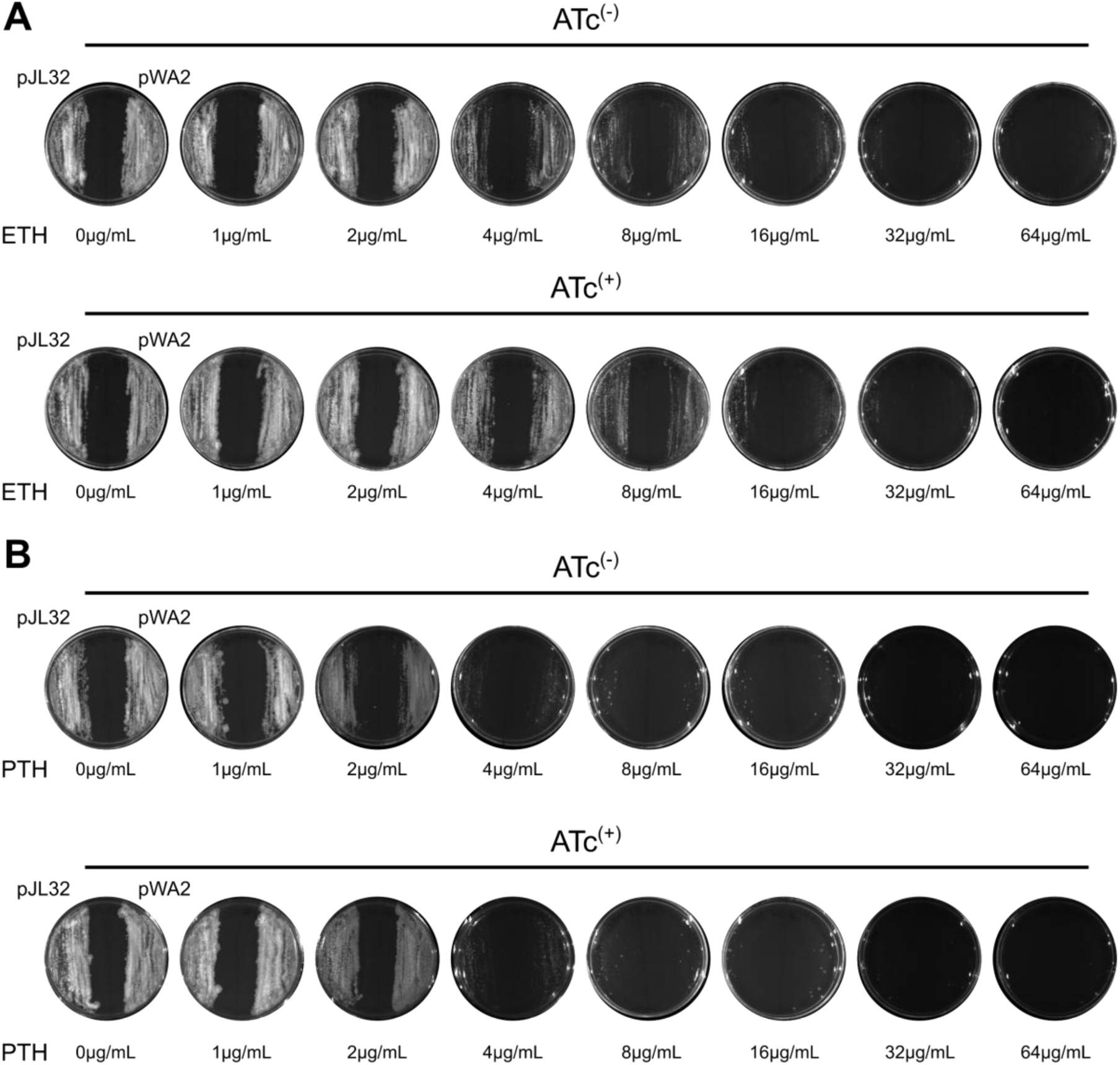
Analysis of cross resistance to ETH and PTH upon *katG* silencing A-B. CRISPRi-targeting of *katG* expression in *Msmeg* does not cause ETH or PTH resistance *in vitro* when assessed on agar plates. Approximately, 10^6^ CFU of exponentially growing *Msmeg* recombinant strains pJL32 or pWA2, were spotted and further evenly spread with a sterile loop on their corresponding quadrant of 7H10 agar plates containing 50 µg·mL⁻¹ of kanamycin, and increasing concentrations (ranging from 0 to 64 µg·mL⁻¹) of ETH **(A)** or PTH **(B)**. Plates were done in the absence of inducer (ATc(−) - top panel) or in the presence of 100 ng·mL⁻¹ of inducer (ATc(+) - bottom panel). For clarity, only bright light acquisitions of the plates are displayed. Results are representative of two biological replicates.

**Figure S3.**
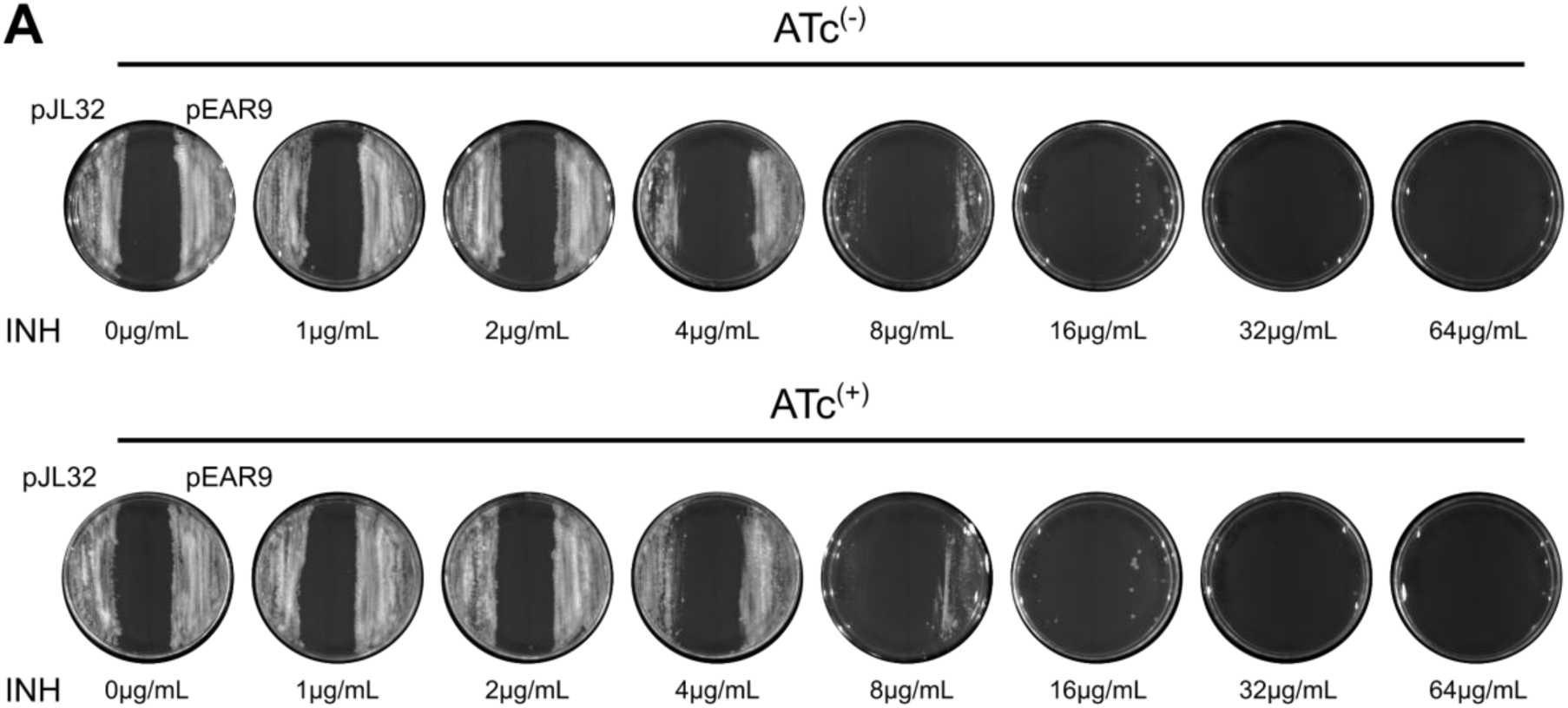
Analysis of cross resistance to INH upon *ethA* silencing. **A-B.** CRISPRi-targeting of *ethA* expression in *Msmeg* does not cause INH resistance *in vitro* when assessed on agar plates. Approximately, 10^6^ CFU of exponentially growing *Msmeg* recombinant strains pJL32 or pEAR9, were spotted and further evenly spread with a sterile loop on their corresponding quadrant of 7H10 agar plates containing 50 µg·mL⁻¹ of kanamycin, and increasing concentrations (ranging from 0 to 64 µg·mL⁻¹) of INH. Plates were done in the absence of inducer (ATc(−) - top panel) or in the presence of 100 ng·mL⁻¹ of inducer (ATc(+) - bottom panel). For clarity, only bright light acquisitions of the plates are displayed. Results are representative of two biological replicates.

**Figure S4.**
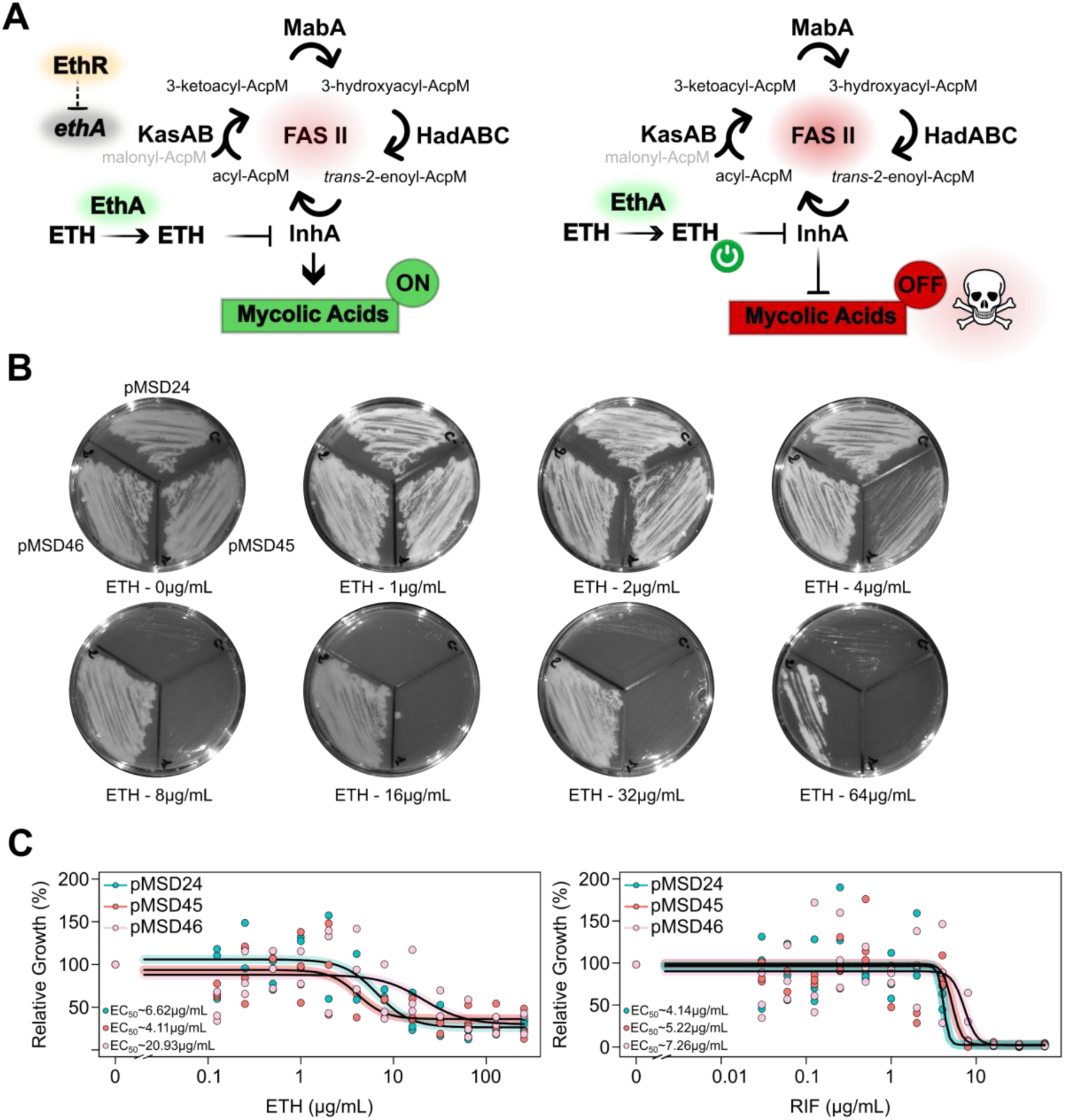
Analysis of *ethA* and *ethR* overexpression on ETH efficacy. **A.** Schematic representation of *ethA* and *ethR* overexpression and their potential effect on the level of mature meromycolic acid required for cell-wall biosynthesis and maintenance upon ETH pulse. Accordingly, genetic overexpression of *ethA* or *ethR* should alter the bioactivation of ETH in its active form and therefore change antibacterial activity. **B.** Overexpression of *ethR,* but not *ethA,* impact *Msmeg* vulnerability towards ETH *in vitro* when assessed on agar plates. Approximately, 10^6^ CFU of exponentially growing *Msmeg* recombinant strains pMS24, pMSD45 and pMSD46 were spotted and further evenly spread with a sterile loop on their corresponding quadrant of 7H10 agar plates containing 50 µg·mL⁻¹ of kanamycin, and increasing concentrations (ranging from 0 to 64 µg·mL⁻¹) of ETH. Results are representative of two biological replicates. **C.** Overexpression of *ethR,* but not *ethA,* impact *Msmeg* vulnerability towards ETH *in vitro* when assessed in broth. Comparative analysis of ETH-mediated growth inhibition against *Msmeg* pMS24 (cyan), pMSD45 (red) and pMSD46 (pink) in Middlebrook 7H9 medium. A control experiment was performed in the presence of rifampicin (RIF). Dose-response analysis displayed were obtained following a 4-parameter nonlinear logistic regression and EC_50_ were determined accordingly. Results are representative of two biological replicates.

**Figure S5.**
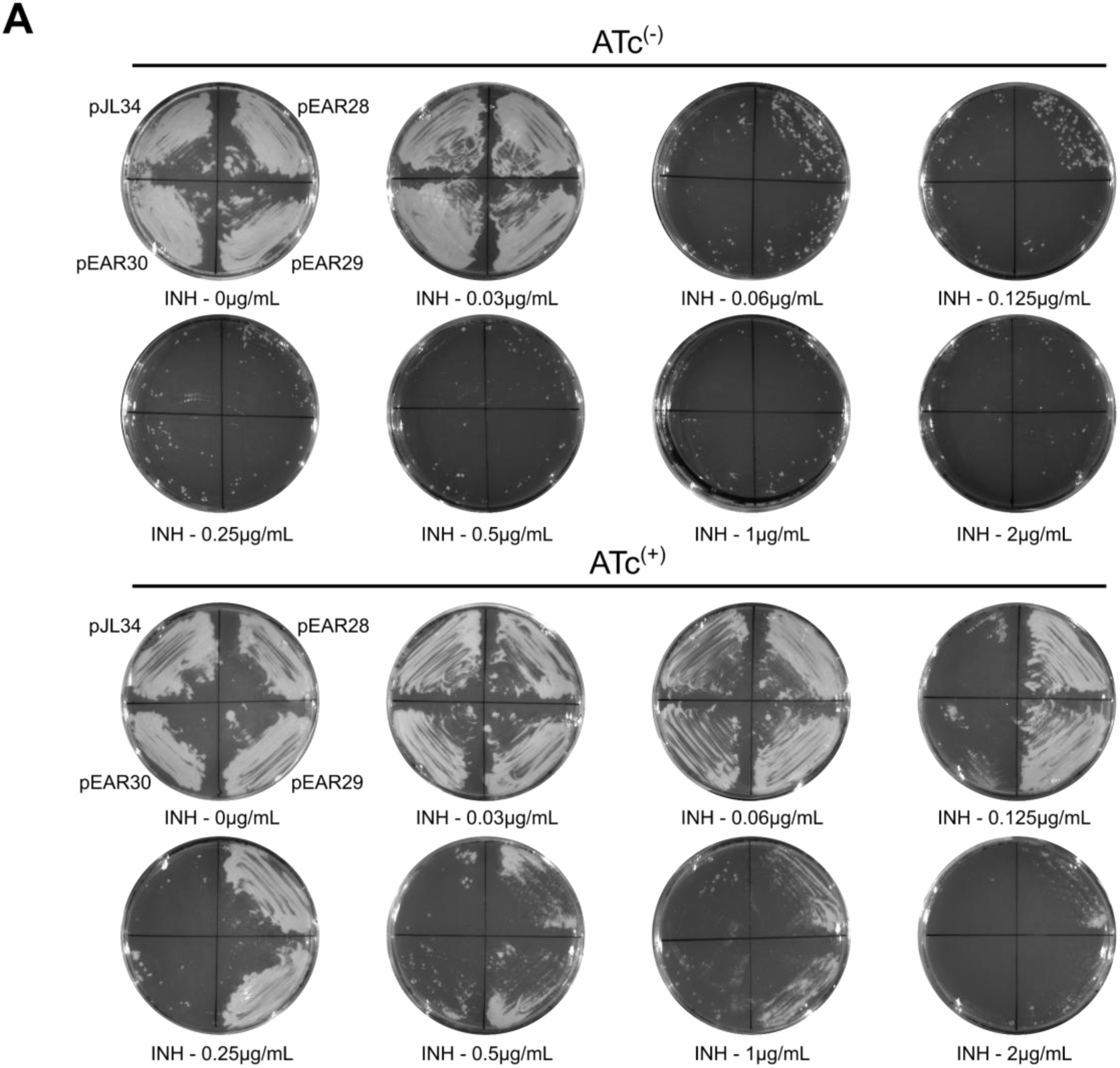
CRISPRi-targeting of *katG* causes conditional resistance to INH in *Mtb.* **A.** CRISPRi-targeting of *katG* expression in *Mtb* causes INH resistance *in vitro* when assessed on agar plates. Approximately, 10^5^ CFU of exponentially growing *Mtb* recombinant strains pJL34, pEAR28, pEAR29 and pEAR30, were spotted and further evenly spread with a sterile loop on their corresponding quadrant of 7H10 OADC agar plates containing 50 µg·mL⁻¹ of kanamycin, and increasing concentrations (ranging from 0 to 2 µg·mL⁻¹) of INH. Plates were done in the absence of inducer (ATc^(−)^ -top panel) or in the presence of 100 ng·mL⁻¹ of inducer (ATc^(+)^ -bottom panel). For clarity, only bright light acquisitions of the plates are displayed. Results are representative of two biological replicates.

**Figure S6.**
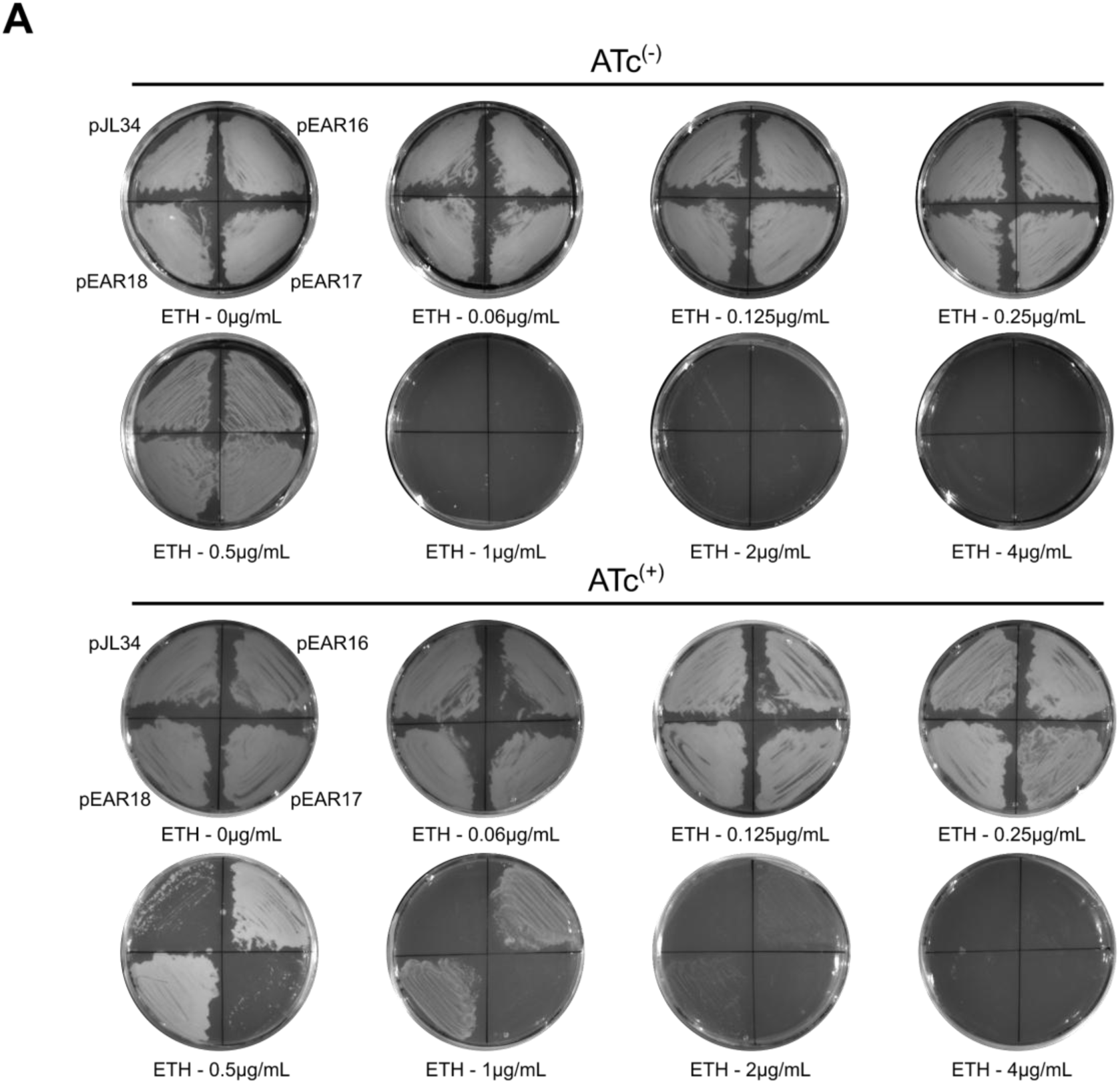
CRISPRi-targeting of *ethA* causes conditional resistance to ETH in *Mtb.* **A.** CRISPRi-targeting of *ethA* expression in *Mtb* causes ETH resistance *in vitro* when assessed on agar plates. Approximately, 10^6^ CFU of exponentially growing *Mtb* recombinant strains pJL34, pEAR16, pEAR17 and pEAR18, were spotted and further evenly spread with a sterile loop on their corresponding quadrant of 7H10 OADC agar plates containing 50 µg·mL⁻¹ of kanamycin, and increasing concentrations (ranging from 0 to 4 µg·mL⁻¹) of ETH. Plates were done in the absence of inducer (ATc^(−)^ -top panel) or in the presence of 100 ng·mL⁻¹ of inducer (ATc^(+)^ -bottom panel). For clarity, only bright light acquisitions of the plates are displayed. Results are representative of two biological replicates.

